# Immune-mediated genetic pathways resulting in pulmonary function impairment increase lung cancer susceptibility

**DOI:** 10.1101/635318

**Authors:** Linda Kachuri, Mattias Johansson, Sara R. Rashkin, Rebecca E. Graff, Yohan Bossé, Venkata Manem, Neil E. Caporaso, Maria Teresa Landi, David C. Christiani, Paolo Vineis, Geoffrey Liu, Ghislaine Scelo, David Zaridze, Sanjay S. Shete, Demetrius Albanes, Melinda C. Aldrich, Adonina Tardón, Gad Rennert, Chu Chen, Gary E. Goodman, Jennifer A. Doherty, Heike Bickeböller, John K. Field, Michael P. Davies, M. Dawn Teare, Lambertus A. Kiemeney, Stig E. Bojesen, Aage Haugen, Shanbeh Zienolddiny, Stephen Lam, Loïc Le Marchand, Iona Cheng, Matthew B. Schabath, Eric J. Duell, Angeline S. Andrew, Jonas Manjer, Philip Lazarus, Susanne Arnold, James D. McKay, Nima C. Emami, Matthew T. Warkentin, Yonathan Brhane, Ma’en Obeidat, Richard M. Martin, Caroline Relton, George Davey Smith, Philip C. Haycock, Christopher I. Amos, Paul Brennan, John S. Witte, Rayjean J. Hung

## Abstract

Impaired lung function is often caused by cigarette smoking, making it challenging to disentangle its role in lung cancer susceptibility. Investigation of the shared genetic basis of these phenotypes in the UK Biobank and International Lung Cancer Consortium (29,266 cases, 56,450 controls) shows that lung cancer is genetically correlated with reduced forced expiratory volume in one second (FEV_1_: *r*_g_=0.098, p=2.3×10^−8^) and the ratio of FEV_1_ to forced vital capacity (FEV_1_/FVC: *r*_g_=0.137, p=2.0×10^−12^). Mendelian randomization analyses demonstrate that reduced FEV_1_ increases squamous cell carcinoma risk (odds ratio (OR)=1.51, 95% confidence intervals: 1.21-1.88), while reduced FEV_1_/FVC increases the risk of adenocarcinoma (OR=1.17, 1.01-1.35) and lung cancer in never smokers (OR=1.56, 1.05-2.30). These findings support a causal role of pulmonary impairment in lung cancer etiology. Integrative analyses reveal that pulmonary function instruments, including 73 novel variants, influence lung tissue gene expression and implicate immune-related pathways in mediating the observed effects on lung carcinogenesis.

## INTRODUCTION

Lung cancer is the most commonly diagnosed cancer worldwide and the leading cause of cancer mortality^1^. Although tobacco smoking remains the predominant risk factor for lung cancer, clinical observations and epidemiological studies have consistently shown that individuals with airflow limitation, particularly those with chronic obstructive pulmonary disease (COPD), have a significantly higher risk of developing lung cancer^2, 3, 4, 5, 6, 7^. Several lines of evidence suggest that biological processes resulting in pulmonary impairment warrant consideration as independent lung cancer risk factors, including observations that previous lung diseases influence lung cancer risk independently of tobacco use^6, 8, 9, 10^, and overlap in genetic susceptibility loci for lung cancer and chronic obstructive pulmonary disease (COPD) on 4q24 (*FAM13A*), 4q31 (*HHIP*), 5q.32 (*HTR4*), the 6p21 region, and 15q25 (*CHRNA3*/*CHRNA5*)^11, 12, 13, 14^. Inflammation and oxidative stress have been proposed as key mechanisms promoting lung carcinogenesis in individuals affected by COPD or other non-neoplastic lung pathologies^9, 11, 15^.

Despite an accumulation of observational findings, previous epidemiological studies have been unable to conclusively establish a causal link between indicators of impaired pulmonary function and lung cancer risk due to the interrelated nature of these conditions^7^. Lung cancer and obstructive pulmonary disease share multiple etiological factors, such as cigarette smoking, occupational inhalation hazards, and air pollution, and 50-70% of lung cancer patients present with co-existing COPD or airflow obstruction^6^. Furthermore, reverse causality remains a concern since pulmonary symptoms may be early manifestations of lung cancer or acquired lung diseases in patients whose immune system has already been compromised by undiagnosed cancer.

Disentangling the role of pulmonary impairment in lung cancer development is important from an etiological perspective, for refining disease susceptibility mechanisms, and for informing precision prevention and risk stratification strategies. In this study we comprehensively assess the shared genetic basis of impaired lung function and lung cancer risk by conducting genome-wide association analyses in the UK Biobank cohort to identify genetic determinants of three pulmonary phenotypes, forced expiratory volume in 1 second (FEV_1_), forced vital capacity (FVC), and FEV_1_/FVC. We examine the genetic correlation between pulmonary function phenotypes and lung cancer, followed by Mendelian randomization (MR) using novel genetic instruments to formally test the causal relevance of impaired pulmonary function, using the largest available dataset of 29,266 lung cancer cases and 56,450 controls from the OncoArray lung cancer collaboration^16^.

## RESULTS

### Heritability and Genetic Correlation

Array-based, or narrow-sense, heritability (*h*_g_) estimates for all lung phenotypes were obtained using LD score regression^17^ based on summary statistics from our GWAS of the UKB cohort (n=372,750 for FEV_1_, n=370,638 for FVC, n=368,817 for FEV_1_/FVC; Supplementary Figure 1) are presented in Table 1. Heritability estimates based on UKB-specific LD scores (n=7,567,036 variants), were consistently lower but more precise than those based on the 1000 Genomes (1000 G) Phase 3 reference population (n=1,095,408 variants). For FEV_1_, *h*_g_ = 0.163 (SE=0.006) and *h*_g_ = 0.201 (SE=0.008), based on UKB and 1000G LD scores, respectively. Estimates for FVC were *h*_g_ = 0.175 (SE=0.007) and *h*_g_ = 0.214 (SE=0.010). Heritability was lower for FEV_1_/FVC: *h*_g_ = 0.128 (SE=0.006) and 0.157 (SE=0.010), based on internal and 1000G reference panels, respectively. For all phenotypes, *h*_g_ did not differ by smoking status and estimates were not affected by excluding the major histocompatibility complex (MHC) region.

**Table 1:** Array-based heritability for FEV_1_, FVC, and FEV_1_/FVC. Estimates were obtained using LD score regression applied to genome-wide summary statistics from the UK Biobank (UKB). Two types of LD scores were used: LD scores estimated using UK Biobank (internal reference population) and pre-computed LD scores based on the 1000 Genomes Phase 3 reference population.

| UKB LD Scores | FEV<sub>1</sub> |  | FVC |  | FEV<sub>1</sub>/FVC |  |
| --- | --- | --- | --- | --- | --- | --- |
|  | h<sub>g</sub> | (SE) | h<sub>g</sub> | (SE) | h<sub>g</sub> | (SE) |
| Overall | 0.163 | (0.006) | 0.175 | (0.007) | 0.128 | (0.006) |
| Never smokers | 0.163 | (0.007) | 0.169 | (0.007) | 0.126 | (0.008) |
| Smokers | 0.159 | (0.007) | 0.172 | (0.009) | 0.129 | (0.008) |
| Overall no MHC | 0.162 | (0.006) | 0.175 | (0.007) | 0.125 | (0.006) |
| 1000G LD Scores |  |  |  |  |  |  |
| Overall | 0.201 | (0.008) | 0.214 | (0.010) | 0.157 | (0.010) |
| Never smokers | 0.209 | (0.010) | 0.215 | (0.011) | 0.159 | (0.011) |
| Smokers | 0.208 | (0.010) | 0.221 | (0.011) | 0.166 | (0.010) |

Partitioning heritability by functional annotation identified large and statistically significant (p<8.5×10^−4^) enrichments for multiple categories (Figure 1; Supplementary Tables 1-3). A total of 35 categories, corresponding to 22 distinct annotations, were significantly enriched for all three pulmonary phenotypes, including annotations that were not previously reported^18^. Large enrichment, defined as the proportion of heritability accounted for by a specific category relative to the proportion of SNPs in that category, was observed for elements conserved in primates^19, 20^ (17.6% of SNPs, 54.7-58.5% of *h*_g_), McVicker background selection statistic^21, 22^ (17.8% of SNPs, 22.6-25.1% of *h*_g_), flanking bivalent transcription starting sites (TSS)/enhancers from Roadmap^20, 23^ (1.4% of SNPs, 11.1-13.2% of *h*_g_), and super enhancers (16.7% of SNPs, 33.9-38.6% of *h*_g_). We also replicated previously reported significant enrichments for histone methylation and acetylation marks H3K4me1, H3K9Ac, and H3K27Ac^18, 24^.

**Figure 1:**
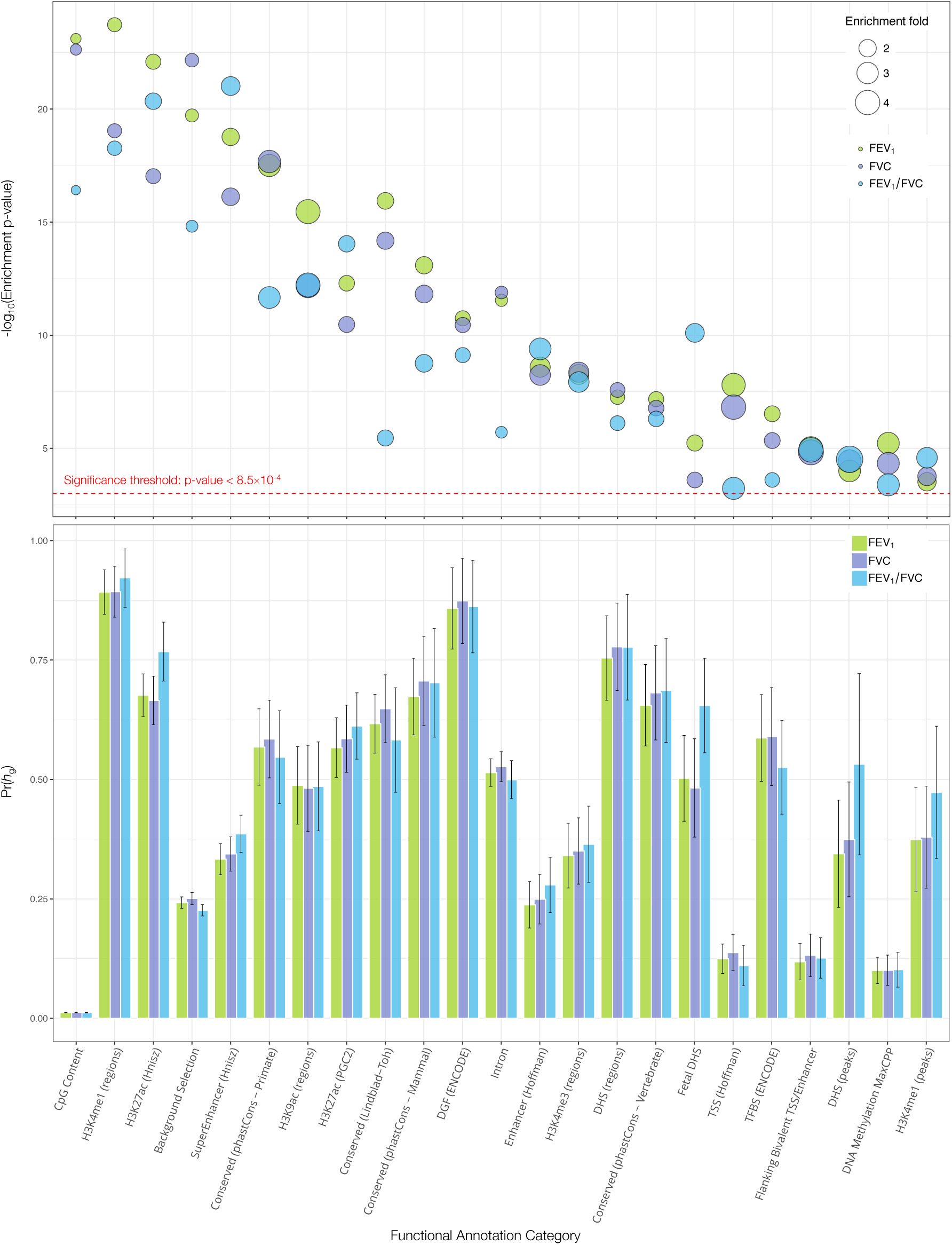
Functional partitioning of array-based heritability for each pulmonary function phenotype. The upper panel depicts the magnitude of category-specific enrichment and corresponding -log_10_(p-value) for 22 distinct functional annotations that were significantly enriched for all three phenotypes (FEV_1_, FVC, FEV_1_/FVC). The lower panel shows the proportion of heritability, Pr(*h*_g_), accounted for by each functional annotation with corresponding standard errors. Functional annotation categories are not mutually exclusive.

Substantial genetic correlation was observed for pulmonary phenotypes with body composition and smoking traits, mirroring phenotypic correlations in epidemiologic studies (Figure 2). Large positive correlations with height were observed for FEV_1_ (*r*_g_=0.568, p=2.5×10^−567^) and FVC (*r*_g_=0.652, p=1.8×10^−864^). Higher adiposity was negatively correlated with FEV_1_ (BMI: *r*_g_=-0.216, p=4.2×10^−74^; percent body fat: *r*_g_=-0.221, p=1.7×10^−66^), FVC (BMI: *r*_g_=-0.262, p=1.6×10^−114^; percent body fat: *r*_g_=-0.254, p=1.2×10^−88^). Smoking status (ever vs. never) was significantly correlated with all lung function phenotypes (FEV_1_ *r*_g_=-0.221, p=8.1×10^−78^; FVC *r*_g_=-0.091, 1.0×10^−16^; FEV_1_/FVC *r*_g_=-0.360, p=7.5×10^−130^). Cigarette pack years and impaired lung function in smokers were also significantly genetically correlated with FEV_1_ (*r*_g_=-0.287 p=1.1×10^−35^), FVC (*r*_g_=-0.253, p=1.9×10^−30^), and FEV_1_/FVC (*r*_g_=-0.108, p=3.0×10^−4^). As a positive control, we verified that FEV_1_ and FVC were genetically correlated with each other (*r*_g_=0.922) and with FEV_1_/FVC (FEV_1_: *r*_g_=0.232 p=4.1×10^−32^; FVC: *r*_g_= -0.167 p=1.0×10^−19^).

**Figure 2:**
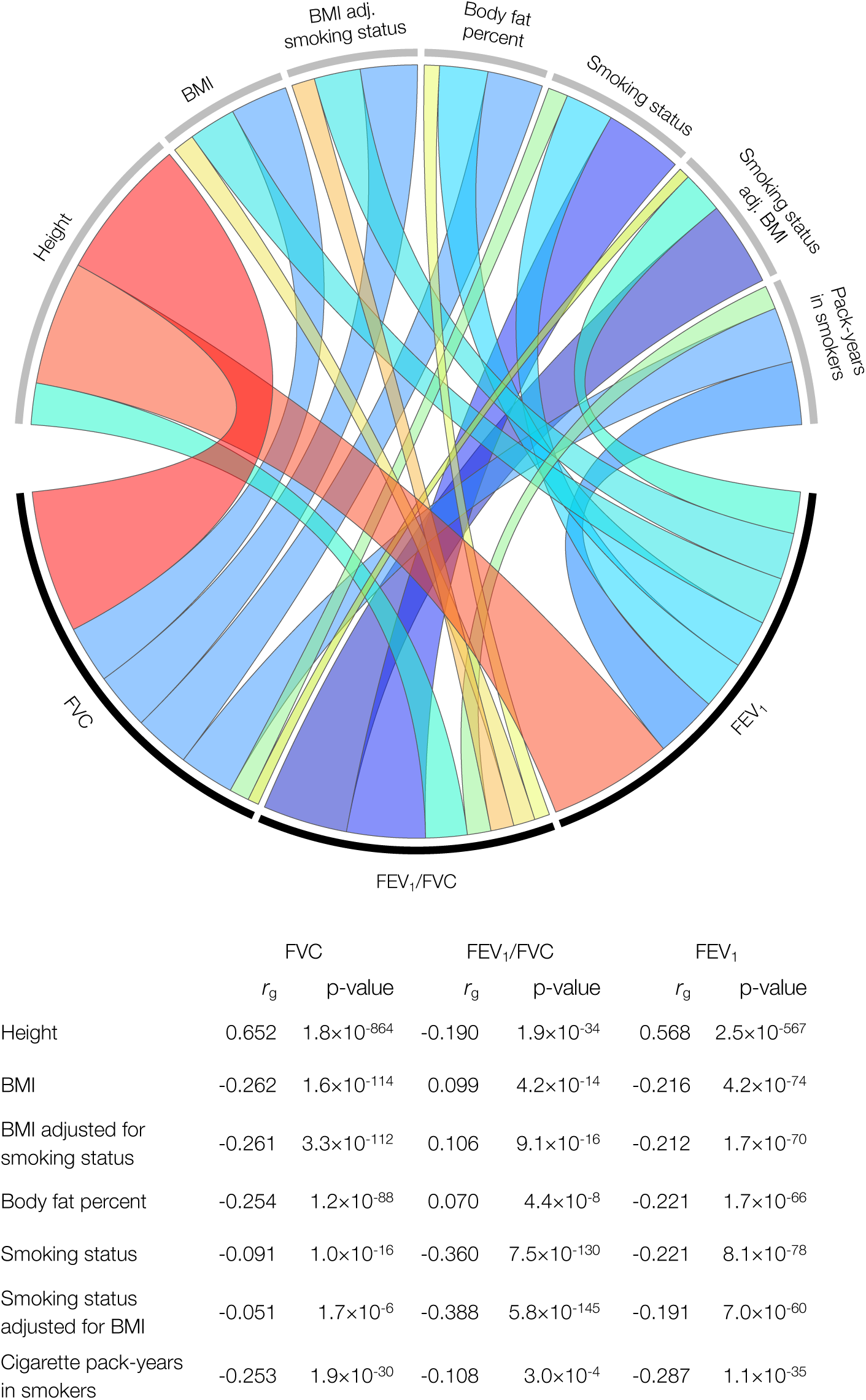
Genetic correlation (*r*_g_) between pulmonary function phenotypes and anthropometric and smoking phenotypes in the UK Biobank cohort. Estimates for *r*_g_ are based on UK Biobank-specific LD scores. The colors in the Circos plot correspond to the direction of genetic correlation, with warm shades denoting positive relationships and cool tones depicting negative correlations. The width of each band in the circos plot is proportional to the magnitude of the absolute value of the *r*_g_ estimate.

Genetic correlations between lung function phenotypes and lung cancer are presented in Figure 3. For simplicity of interpretation coefficients were rescaled to represent genetic correlation with impaired (decreasing) lung function. Impaired FEV_1_ was positively correlated with lung cancer overall (*r*_g_=0.098, p=2.3×10^−8^), squamous cell carcinoma (*r*_g_=0.137, p=7.6×10^−9^), and lung cancer in smokers (*r*_g_=0.140, p=1.2×10^−7^). Genetic correlations were attenuated for adenocarcinoma histology (*r*_g_=0.041, p=0.044) and null for never-smokers (*r*_g_=-0.002, p=0.96). A similar pattern of associations was observed for FVC. Reduced FEV_1_/FVC was positively correlated with all lung cancer subgroups (overall: *r*_g_=0.137, p=2.0×10^−12^; squamous carcinoma: *r*_g_=0.137, p=4.3×10^−8^; adenocarcinoma: *r*_g_=0.125, p=7.2×10^−9^; smokers: *r*_g_=0.185, p=1.4×10^−10^), except for never smokers (*r*_g_=0.031, p=0.51).

**Figure 3:**
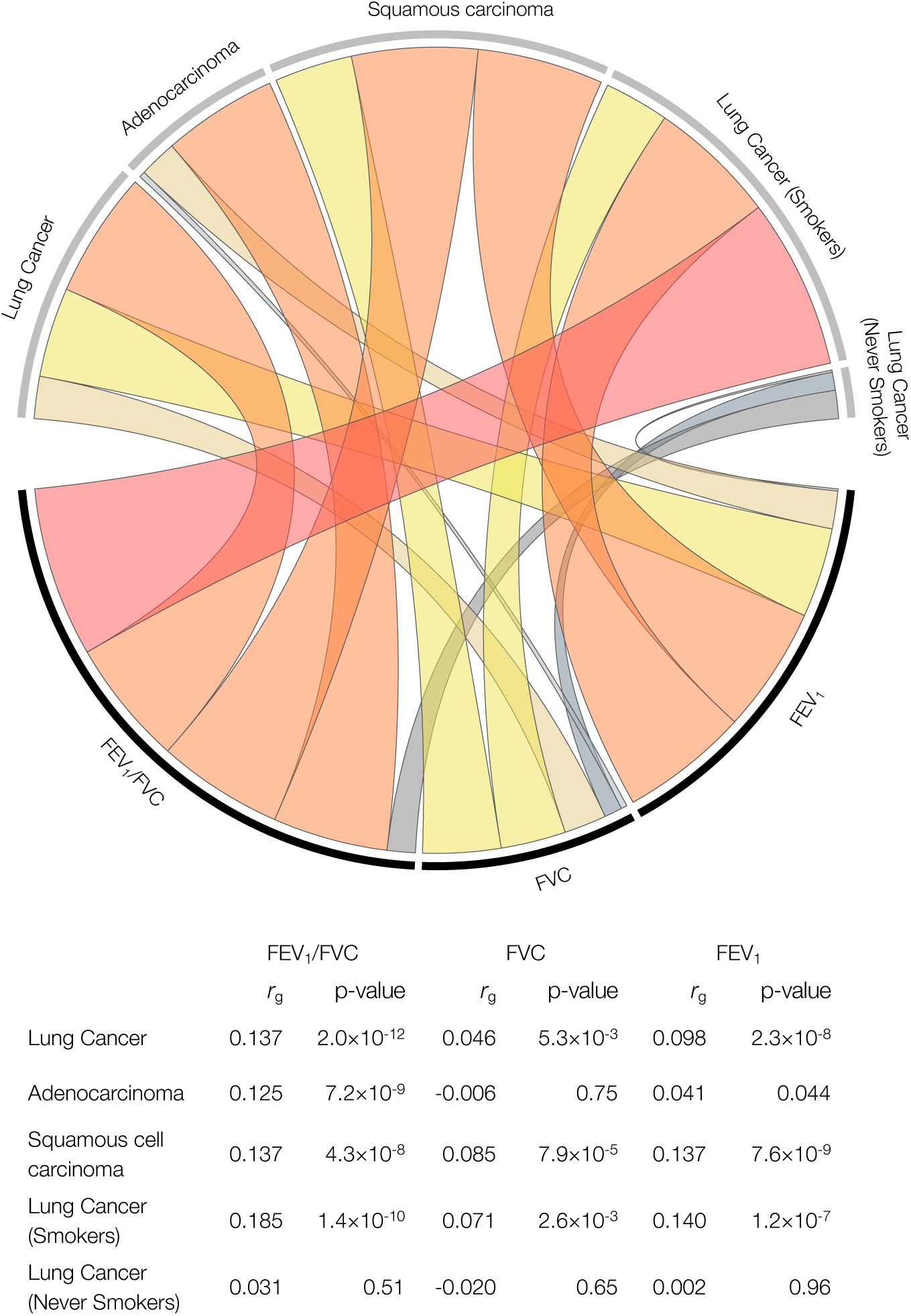
Genetic correlation (*r*_g_) between pulmonary function phenotypes and lung cancer subtypes. Estimates of *r*_g_ are based on genome-wide summary statistics from the UK Biobank cohort for pulmonary traits, and the International Lung Cancer Consortium OncoArray study for lung cancer. Genetic correlations have been re-scaled to depict associations between impaired (reduced) pulmonary function and lung cancer risk. The colors in the circos plot correspond to the direction of genetic correlation, with warm shades depicting positive correlations between impaired pulmonary function and lung cancer risk, and gray tones corresponding to inverse and null correlations. The width of each band in the circos plot is proportional to the magnitude of the absolute value of the *r*_g_ estimate.

Exploring the functional underpinnings of these genetic correlations revealed three functional categories that were significantly enriched for lung cancer (Supplementary Table 4), and have not been previously reported^25^. All these categories were also significantly enriched for pulmonary traits. CpG dinucleotide content^22^ included only 1% of SNPs, but had a strong enrichment signal for lung cancer (p=2.1×10^−7^), FEV_1_ (p=7.7×10^−24^), FVC (p=2.3×10^−23^, and FEV_1_/FVC (p=3.8×10^−17^). Other shared features included background selection (lung cancer: p=1.0×10^−6^, FEV_1_: p=1.9×10^−20^, FVC: p=6.9×10^−23^, FEV_1_/FVC: p=1.5×10^−15^) and super enhancers (lung cancer: p=4.4×10^−6^, FEV_1_: p=3.4×10^−24^, FVC: p=5.1×10^−20^, FEV_1_/FVC: p=9.6×10^−22^).

### Genome-Wide Association Analysis for Instrument Development

Based on the results of our GWAS in the UK Biobank, we identified 207 independent instruments for FEV_1_ (*P*<5×10^−8^, replication *P*<0.05; LD *r*^2^<0.05 within 10,000 kb), 162 for FVC, and 297 for FEV_1_/FVC. We confirmed that our findings were not affected by spirometry performance quality, with a nearly perfect correlation between effect sizes (*r*^2^=0.995, p=2.5×10^−196^) in the main discovery analysis and after excluding individuals with potential blow acceptability issues (Field 3061≠0; n=60,299). After applying these variants to the lung cancer OncoArray dataset and selecting LD proxies (*r*^2^>0.90) for unavailable variants, the final set of instruments consisted of 193 variants for FEV_1_, 144 for FVC, and 264 SNPs for FEV_1_/FVC (Supplementary Data 1-3), for a total of 601 instruments. The proportion of trait variation accounted for by each set of instruments was estimated in the UKB replication sample consisting of over 110,00 individuals (Supplementary Figure 1), and corresponded to 3.13% for FEV_1_, 2.27% for FVC, and 5.83% for FEV_1_/FVC. We also developed instruments specifically for never smokers based on a separate GWAS of this population, which yielded 76 instruments for FEV_1_, 112 for FEV_1_/FVC, and 57 for FVC, accounting for 2.06%, 4.21%, and 1.36% of phenotype variation, respectively (Supplementary Data 4-6).

After removing overlapping instruments between pulmonary phenotypes and LD-filtering (*r*^2^<0.05) across the three traits, 447 of the 601 variants were associated with at least one of FEV_1_, FVC, or FEV_1_/FVC (*P*<5×10^−8^, replication *P*<0.05). We compared these 447 independent variants to the 279 lung function variants recently reported by Shrine et al.^18^ based on an analysis of the UK Biobank and SpiroMeta consortium, by performing clumping with respect to these index variants (LD *r*^2^<0.05 within 10,000 kb). Our set of instruments included an additional 73 independent variants, 69 outside the MHC region (Supplementary Table 5), that achieved replication at the Bonferroni-corrected threshold for each trait (maximum replication *P* = 2.0×10^−4^).

Our instruments included additional independent signals in known lung function loci and variants in genes newly linked to lung function, such as *HORMAD2* at 22q12.1 (rs6006399: *P*_FEV1_=1.9×10^−18^), which is involved in synapsis surveillance in meiotic prophase, and *RIPOR1* at 16q22.1 (rs7196853: *P* _FEV1/FVC_=1.3×10^−16^), which plays a role in cell polarity and directional migration. Several new variants further support the importance of the tumor growth factor beta (TGF-β) signaling pathway, including *CRIM1* (rs1179500: *P*_FEV1/FVC_=3.6×10^−17^) and *FGF18* (rs11745375: *P*_FEV1/FVC_=1.6×10^−11^). Another novel gene, *PIEZO1* (rs750739: *P*_FEV1_=1.8×10^−10^), encodes a mechano-sensory ion channel, supports adaptation to stretch of the lung epithelium and endothelium, and promotes repair after alveolar injury^26, 27^. In never smokers a signal was identified at 6q15 in *BACH2* (rs58453446: *P*_FEV1/FVC-nvsmk_=8.9×10^−10^), a gene required for pulmonary surfactant homeostasis. Lastly, two lung function variants mapped to genes somatically mutated in lung cancer: *EML4* (rs12466981: *P*_FEV1/FVC_=2.7×10^−14^) and *BRAF* (rs13227429: *P*_FVC_=5.6×10^−9^).

### Mendelian Randomization

The causal relevance of impaired pulmonary function was investigated by applying genetic instruments developed in the UK Biobank to the OncoArray lung cancer dataset, comprised of 29,266 lung cancer cases and 56,450 controls (Supplementary Table 6). Primary analyses were based on the maximum likelihood (ML) and inverse variance weighted (IVW) multiplicative random effects estimators^28, 29^. Sensitivity analyses were conducted using the weighted median (WM) and robust adjusted profile score (RAPS) estimators^30,31^. A genetically predicted decrease in FEV_1_ was significantly associated with increased risk of lung cancer overall (OR_ML_=1.28, 95% CI: 1.12-1.47, p=3.4×10^−4^) and squamous carcinoma (OR_ML_=2.04, 1.64-2.54, p=1.2×10^−10^), but not adenocarcinoma (OR_ML_=0.99, 0.83-1.19, p=0.96) (Figure 4; Supplementary Table 7). The association with lung cancer was not significant across all estimators (OR_WM_=1.06, p=0.57; OR_RAPS_=1.13, p=0.26). There was no evidence of directional pleiotropy based on the MR Egger intercept test (β_0 *Egger*_ ≠0, p<0.05), but significant heterogeneity among SNP-specific causal effect estimates was observed, which may be indicative of balanced horizontal pleiotropy (lung cancer: *P*_Q_=2.1×10^−41^; adenocarcinoma: *P*_Q_=3.4×10^−9^; squamous carcinoma: *P*_Q_=1.1×10^−30^). After excluding outlier variants contributing to this heterogeneity, 36 for lung cancer and 34 for squamous carcinoma, the association with FEV_1_ diminished for both phenotypes (lung cancer: OR_ML_=OR_IVW_=1.12, p=0.13), but remained statistically significant for squamous carcinoma (OR_IVW_=1.51, 1.21-1.88, p=2.2×10^−4^), with comparable effects observed using other estimators (OR_ML_=1.50, p=6.7×10^−4^; ORRAPS=1.48, p=1.7×10^−3^; OR_WM_=1.44, p=0.040).

**Figure 4:**
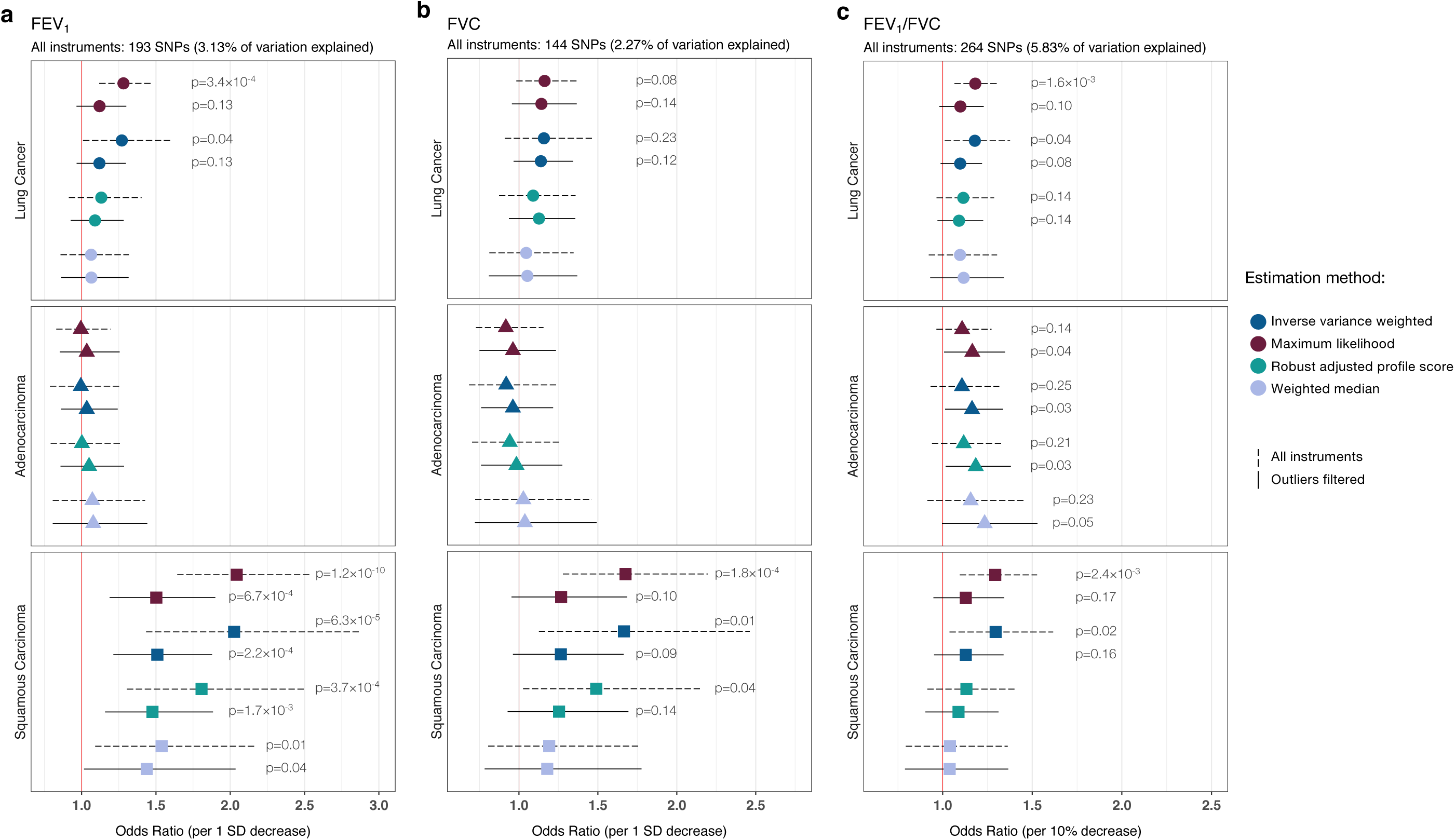
Odds ratios (OR) and 95% confidence intervals for the effect of impaired pulmonary function on lung cancer risk, estimated using Mendelian randomization (MR). Multiple MR estimation methods were applied to the International Lung Cancer Consortium OncoArray dataset, comprised of 29,266 lung cancer cases (11,273 adenocarcinoma, 7,426 squamous cell carcinoma) and 56,450 controls to assess the causal relevance of impaired FEV_1_ **(a)**, FVC **(b)**, and FEV_1_/FVC **(c)**. MR estimates based on the full set of genetic instruments are compared to estimates after excluding outliers suspected of violating MR assumptions. Only associations with p-values less than 0.25 are labeled. Proportion of variation explained by the genetic instruments was estimated in a separate replication sample of over 110,000 individuals from the UK Biobank.

Genetic predisposition to reduced FVC was inconsistently associated with squamous carcinoma risk (OR_ML_=1.68, p=1.8×10^−4^; OR_WM_=1.19, p=0.38). Effects became attenuated and more similar after removing outliers (OR_ML_=1.27, p=0.10; OR_RAPS_=1.25, p=0.14) (Figure 4; Supplementary Table 8). A genetically predicted 10% decrease in FEV_1_/FVC was associated with an elevated risk of lung cancer in some models (OR_ML_=1.18, 1.07-1.31, p=1.6×10^−3^), but not others (OR_WM_=1.10, p=0.30; OR_RAPS_=1.11, p=0.14) (Figure 4; Supplementary Table 9). The association with squamous carcinoma was also inconsistent across estimators. After removing outliers contributing to significant effect heterogeneity (lung cancer: *P*_Q_=1.2×10^−28^; adenocarcinoma: *P*_Q_=3.4×10^−9^; squamous carcinoma: *P*_Q_=5.3×10^−15^), the association with adenocarcinoma strengthened (OR_ML_=1.17, 1.01-1.35; OR_RAPS_=1.18, 1.02-1.38), while associations for lung cancer and squamous carcinoma became attenuated.

We examined the cancer risk in never smokers, by applying genetic instruments developed specifically in this population, to 2355 cases and 7504 controls (Figure 5; Supplementary Table 10). A genetically predicted 1-SD decrease in FEV_1_ and FVC was not associated with lung cancer risk in never smokers. However, a 10% reduction in FEV_1_/FVC was associated with a 61% increased risk (OR_ML_=1.61, 1.10-2.35, p=0.014; OR_IVW_=1.60, p=0.030). Outlier filtering did not have an appreciable impact on the results (OR_ML_=1.56, 1.05-2.30, p=0.027; OR_IVW_=1.55, 1.05-2.28, p=0.028). A sensitivity analysis applied to 264 FEV_1_/FVC instruments not specific to never smokers yielded an attenuated estimate (OR_IVW_=1.35, 1.03-1.75, p=0.027), but confirmed the impact of FEV_1_/FVC reduction on lung cancer risk.

**Figure 5:**
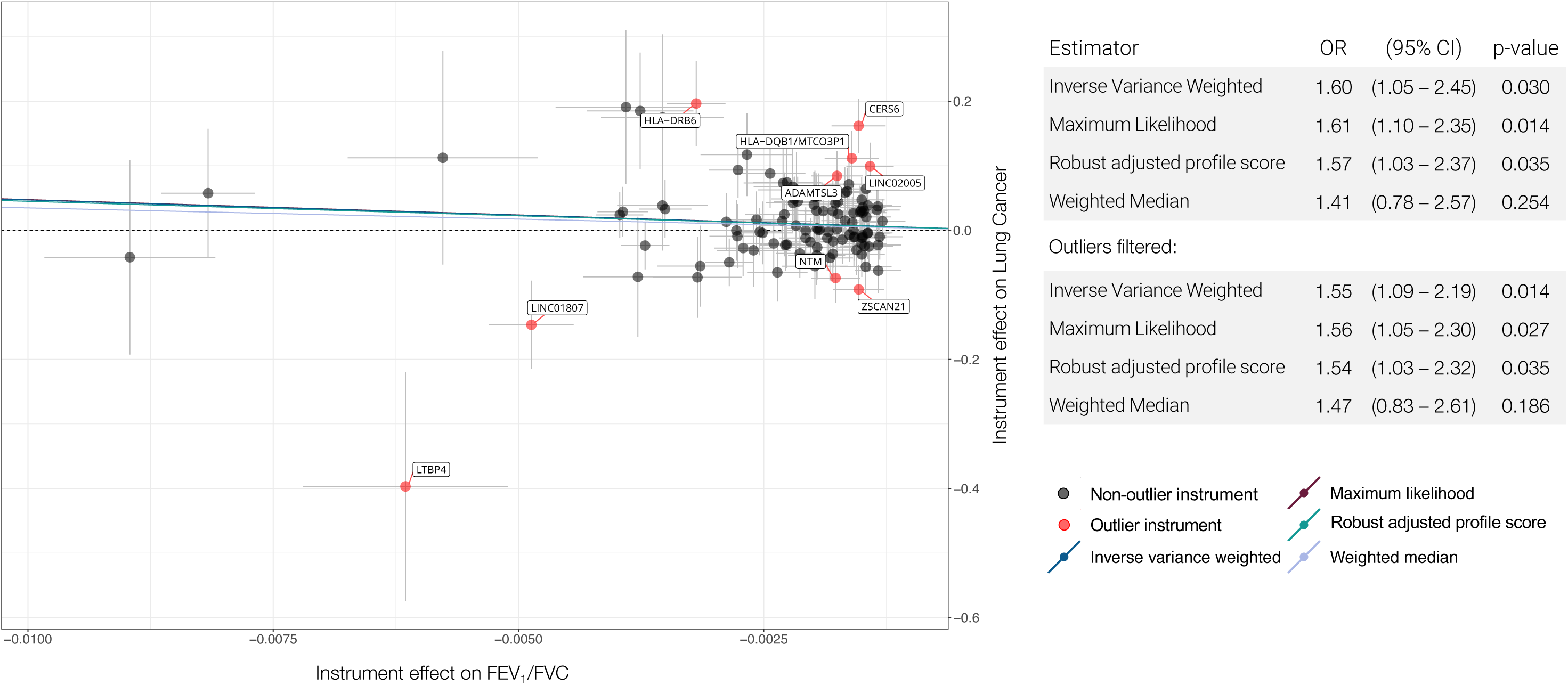
Scatterplot depicting the Mendelian randomization (MR) results for FEV_1_/FVC in never smokers and lung cancer in never smokers (2355 cases, 7504 controls). The scatterplot illustrates the effects of individual instruments on FEV_1_/FVC and lung cancer risk, highlighting potentially invalid outlier instruments that were filtered. Individual instrument effects in the scatterplot correspond to a 1-unit decrease in FEV_1_/FVC, but the summary odds ratios (ORs) for lung cancer have been rescaled to correspond to a 10% decrease in FEV_1_/FVC. Summary log(ORs) based on different MR estimators correspond to the slope of the lines in scatterplot

For completeness, we also present MR estimates for the effect of impaired pulmonary function on lung cancer risk in smokers (Supplementary Table 11). Despite the larger sample size (23,223 cases and 16,964 controls) compared to never smokers, a genetically predicted 10% reduction in FEV_1_/FVC was weakly and inconsistently associated with lung cancer risk (OR_IVW_=1.15, p=0.038; OR_RAPS_=1.08, p=0.488). Genetic predisposition to FEV_1_ and FVC impairment did not appear to confer an increased risk among smokers.

Extensive MR diagnostics are summarized in Supplementary Table 12. All analyses used strong instruments (F-statistic > 40) and did not appear to be weakened by violations of the no measurement error (NOME) assumption (*I*^2^_GX_ statistic >0.97). MR Steiger test^32^ was used to orient the causal effects and confirmed that instruments for pulmonary function were affecting lung cancer susceptibility, not the reverse, and this direction of effect was highly robust. No instruments were removed based on Steiger filtering. We also confirmed that none of the genetic instruments were associated with nicotine dependence phenotypes (*P*<1×10^−5^), such as time to first cigarette, difficulty in quitting smoking, and number of quit attempts, which were available for a subset of individuals in the UKB. All MR analyses were adequately powered, with >80% power to detect a minimum OR of 1.25 for FEV_1_ and FEV_1_/FVC (Supplementary Figure 2). For never smokers, we had 80% power to detect a minimum OR of 1.40 for FEV_1_/FVC and 1.60 for FEV_1_.

Given the genetic correlation observed for pulmonary phenotypes cigarette smoking and adiposity, we conducted several sensitivity analyses to further address any potential confounding by these phenotypes. The finding for squamous carcinoma and FEV_1_ was further interrogated using multivariable MR (MVMR) by incorporating genetic instruments for BMI^33^ and smoking behavior^34^ to estimate the direct effect of FEV_1_ on squamous carcinoma risk. MVMR using all instruments yielded an OR of 1.95 (95% CI: 1.36-2.80, p=2.8×10^−4^) per 1-SD decrease in FEV_1_ and an OR of 1.63 (95% CI: 1.20-2.23, p=1.8×10^−3^) after filtering outlier instruments.

We confirmed that none of the genetic instruments were associated with smoking status (ever/never), cigarette pack-years (continuous), or adiposity (body fat percentage) at the *P* < 5×10^−8^ level. However, several variants were associated based on a *P* < 1×10^−5^ threshold (25 for FEV_1_ and 18 for FEV_1_/FVC). We repeated MR analyses after removing these variants (Supplementary Table 13) and confirmed that our results remained robust for FEV_1_ and squamous cell carcinoma (OR_IVW_=2.02, 1.40-2.92, p=1.9×10^−4^) and FEV_1_/FVC and adenocarcinoma (OR_IVW_=1.19, 1.01-1.40, p=0.04). However, there was still significant heterogeneity among the causal effect estimates. After filtering the remaining outliers, the effect of a 10% decrease in FEV_1_/FVC on adenocarcinoma strengthened (OR_IVW_=1.24, 1.08-1.43, p=2.4×10^−3^), while estimates attenuated slightly for FEV1 and squamous carcinoma (OR_IVW_=1.46, 1.14-1.87, p=2.7×10^−3^).

We also considered the possibility of residual confounding in our GWAS due to insufficient adjustment for smoking-related factors. We thus re-estimated SNP effects on FEV_1_, FVC, and FEV_1_/FVC with adjustment for continuous cigarette pack-years and years since quitting. The distribution of effect sizes did not differ between the two analyses (p>0.05), and the correlation with our original instrument weights was strong for all phenotypes (Pearson’s *r* ≥ 0.87, p<1×10^−40^) (Supplementary Figure 3).

Lastly, we examined the association between FEV_1_ and FEV_1_/FVC genetic instruments and COPD, defined as FEV_1_/FVC<0.70. Among FEV_1_ instruments, 64% (123 variants) were associated with COPD at p<0.05 and 16% (31 variants) at p<5×10^−8^ (Supplementary Figure 4). All instruments for FEV_1_/FVC were associated with COPD at the nominal level, and 40% (105 variants) reached genome-wide significance. Using weights from estimated associations between the 105 instruments and COPD log(OR), we observed a modestly increased risk of lung adenocarcinoma (OR_IVW_=1.08, 1.01-1.15, p=0.015), which parallels our findings based on instruments developed for the continuous FEV_1_/FVC phenotype.

### Functional Characterization of Lung Function Instruments

To gain insight into biological mechanisms mediating the observed effects of impaired pulmonary function on lung cancer risk, we conducted in-silico analyses of functional features associated with the genetic instruments for each lung phenotype.

We identified 185 statistically significant (Bonferroni p<0.05) *cis*-eQTLs for 101 genes among the genetic instruments for FEV_1_ and FEV_1_/FVC based on lung tissue gene expression data from the Laval biobank^35^ (Supplementary Data 7). Predicted expression of 7 genes was significantly (p<5.0×10^−4^) associated with lung cancer risk: *SECISBP2L, HLA-L, DISP2, MAPT, KANSL1-AS1, LRRC37A4P*, and *PLEKHM1* (Supplementary Figure 5). Of these, *SECISBP2L* (OR=0.80, p=5.2×10^−8^), *HLA-L* (OR=0.84, p=1.6×10^−6^), and *DISP2* (OR=1.25, p=1.6×10^−4^) displayed consistent directions of effect for pulmonary function and lung cancer risk, whereby alleles associated with increased expression were associated with impaired FEV_1_ or FEV_1_/FVC and increased cancer risk (or conversely, positively associated with pulmonary function and inversely associated with cancer risk). Gene expression associations with inconsistent effects are more likely to indicate pleiotropic pathways not operating primarily through pulmonary impairment. Differences by histology were observed for *SECISBP2L*, which was associated with adenocarcinoma (OR=0.54, p=3.1×10^−14^), but not squamous cell carcinoma (OR=1.05, p=0.44). Effects observed for *DISP2* (OR=1.21, p=0.021) and *HLA-L* (OR=0.90, p=0.034) were attenuated for adenocarcinoma, but not for squamous carcinoma (*DISP2*: OR=1.30, p=6.2×10^−3^; *HLA-L*: OR=0.75, p=1.6×10^−6^).

A total of 70 lung function instruments were mapped to genome-wide significant (p<5.0×10^−8^) protein quantitative trait loci (pQTL) affecting the plasma levels of 64 different proteins (Supplementary Data 8), based on data from the Human Plasma Proteome Atlas^36^. Many of these pQTL targets are involved in regulation of immune and inflammatory responses, such as interleukins (IL21, IL1R1, IL17RD, IL18R1), MHC class I polypeptide-related sequences, transmembrane glycoproteins expressed by natural killer cells, and members of the tumor necrosis receptor superfamily (TNFSF12, TNFRSF6B, TR19L). Other notable associations include NAD(P)H dehydrogenase [quinone] 1 (NQO1) a detoxification enzyme involved in protecting lung tissues in response to reactive oxidative stress (ROS) and promoting p53 stability^37^. NQO1 is a target of the NFE2-related factor 2 (NRF2), a master regulator of cellular antioxidant response that has generated considerable interest as a chemoprevention target^38, 39^.

Next, we analyzed genes where the lung function instruments were localized using curated pathways from the Reactome database. Significant enrichment (FDR q<0.05) was observed only for FEV_1_/FVC instruments in never smokers, with an over-representation of pathways involved in adaptive immunity and cytokine signaling (Supplementary Figure 6). Top-ranking pathways with q=2.2×10^−6^ included translocation of ZAP-70 to immunological synapse, phosphorylation of CD3 and TCR zeta chains, and PD-1 signaling. These findings are in line with the predominance of immune-related pQTL associations. Examining all instruments for FEV_1_ and FEV_1_/FVC identified significant over-representation (FDR q<0.05) of six immunologic signatures from the ImmuneSigDB collection^40^, including pathways implicated in host response to infection and immunization (Supplementary Figure 7).

## DISCUSSION

Despite a substantial body of observational literature demonstrating an increased risk of lung cancer in individuals with pulmonary dysfunction^2, 3, 4, 5, 6, 7, 41^, confounding by shared environmental risk factors and high co-occurrence of lung cancer and airflow obstruction created uncertainty regarding the causal nature of this relationship. We comprehensively investigated this by characterizing shared genetic profiles between lung cancer and lung function, and interrogated causal hypotheses using Mendelian randomization, which overcomes many limitations of observational studies. We also provide insight into biological pathways underlying the observed associations by incorporating functional annotations into heritability analyses, assessing eQTL and pQTL effects of lung function instruments, and conducting pathway enrichment analyses.

The large sample size of the UK Biobank allowed us to successfully create instruments for three pulmonary function phenotypes, FEV_1_, FEV_1_/FVC, and FVC. Although these phenotypes are closely related, they capture different aspects of pulmonary impairment, with FEV_1_ and FEV_1_/FVC used for diagnostic purposes in clinical setting. Our genetic instruments captured known and novel mechanisms involved in pulmonary function. Of the 73 novel variants identified here, many were in loci implicated in immune-related functions and pathologies. Examples include *HORMAD2*, which has been previously linked to inflammatory bowel disease^42, 43^ and tonsillitis^44^, and *RIPOR1* (also known as *FAM65A*), which is part of a gene expression signature for atopy^45^. *PIEZO1* is primarily involved in mechano-transduction and tissue differentiation during embryonic development^46, 47, 48^, however recent evidence has emerged delineating its role in optimal T-cell receptor activation and immune regulation^49^. *BACH2*, the new signal for FEV_1_/FVC in never smokers, is involved in alveolar macrophage function^50^, as well as selection-mediated *TP53* regulation and checkpoint control^51^. The lead variant identified here is independent (*r*^2^<0.05) of *BACH2* loci nominally associated with lung function decline in a candidate gene study of COPD patients^52^, suggesting there may be differences in the genetic architecture of pulmonary traits in never smokers.

Our genetic correlation analyses indicate shared genetic determinants between pulmonary function with anthropometric traits and cigarette smoking. Our results are in contrast with the recent findings of Wyss et al.^24^, who did not observe statistically significant genetic correlations for any pulmonary function phenotypes with height and smoking, as well FVC and FEV_1_/FVC, using publicly available summary statistics from the UKB and other studies of European ancestry individuals. In this respect, assessing genetic correlation within a single well-characterized population provides improved power while minimizing potential for bias and heterogeneity when combining data from multiple sources.

We observed statistically significant genetic correlations between pulmonary function impairment and lung cancer susceptibility for all lung cancer subtypes, except for never smokers. Reduced FEV_1_/FVC was significantly correlated with increased risk of lung cancer overall, squamous cell carcinoma, and adenocarcinoma. Significant genetic correlations with FEV_1_ and FVC were observed for lung cancer overall, in smokers, and for tumors with squamous cell histology, but not adenocarcinoma. Jiang et al.^25^ reported a similar magnitude of genetic correlation with FEV_1_/FVC, but did not observe an association with FVC, and did not assess FEV_1_. Differences in our results may be attributable to their use of GWAS summary statistics for pulmonary phenotypes from the interim UK Biobank release. Our findings demonstrate substantial overlap in the genetic architecture of obstructive and neoplastic lung disease, particularly for highly conserved variants that are likely to be subject to natural selection, and super enhancers. However, genetic correlations do not support a causal interpretation, especially considering the shared heritability with potentially confounding traits, such as smoking and obesity.

On the other hand, Mendelian randomization analyses revealed histology-specific effects of reduced FEV_1_ and FEV_1_/FVC on lung cancer susceptibility, suggesting that these indicators of impaired pulmonary function may be causal risk factors. Genetic predisposition to FEV_1_ impairment conferred an increased risk of lung cancer overall, particularly for squamous carcinoma. This relationship persisted after filtering potentially pleiotropic instruments and performing other sensitivity analyses, including multivariable Mendelian randomization and manual filtering of variants associated with smoking or adiposity. FEV_1_/FVC reduction appeared to increase the risk of lung adenocarcinoma, as well as lung cancer among never-smokers. The latter finding is particularly compelling since it precludes confounding by smoking-related factors and demonstrates an association with the most clinically relevant pulmonary phenotype. The increased lung cancer risk in never smokers was also observed using genetic instruments developed specifically in never smokers and in sensitivity analyses using instruments from the population that also includes smokers. We hypothesize that the effects of pulmonary obstruction are mediated by chronic inflammation and immune response, which is supported by the overrepresentation of adaptive immunity and cytokine signaling pathways and pQTL effects among FEV_1_ and FEV_1_/FVC instruments.

Examining lung eQTL effects of our genetic instruments identified additional relevant mechanisms, including gene expression of *SECISBP2L* and *DISP2. SECISBP2L* at 15q21 is essential for ciliary function^53^ and has an inhibitory effect on lung tumor growth by suppressing cell proliferation and inactivation of Aurora kinase A^54^. This gene was among several susceptibility regions identified in the most recent lung cancer GWAS^16^, and now we more conclusively establish impaired pulmonary function as the mechanism mediating *SECISBP2L* effects on risk of lung cancer overall, particularly adenocarcinoma. Less is known about *DISP2*, although it has been implicated in the conserved Hedgehog signaling pathway essential for embryonic development and cell differentiation^55^.

One of the main challenges and outstanding questions in previous epidemiologic studies has been clarifying how smoking fits into the causal pathway between impaired pulmonary function and lung cancer risk. Are indicators of airway obstruction simply proxies for smoking-induced carcinogenesis? The association between reduced FEV_1_/FVC and risk of adenocarcinoma and lung cancer in never smokers observed in our Mendelian randomization analysis and in previous studies^8, 9^, argues against this simplistic explanation and points to alternative pathways. Chronic airway inflammation fosters a lung microenvironment with altered signaling pathways, aberrant expression of cytokines, chemokines, growth factors, and DNA damage-promoting agents, all of which promote cancer initaiton^15^. This mechanism may be particularly relevant for adenocarcinoma, which is the most common lung cancer histology in never smokers, arising from the peripheral alveolar epithelium that has less direct contact with inhaled carcinogens.

Dysregulated immune function is a hallmark of lung cancer and COPD, with both diseases sharing similar inflammatory cell profiles characterized by macrophages, neutrophils, and CD4+ and CD8+ lymphocytes. Immune cells in COPD and emphysema exhibit T helper 1 (Th1)/Th17 polarization, decreased programmed death ligand-1 (PD-L1) expression in alveolar macrophages, and increased production of interferon (IFN)-γ by CD8+ T cells^56^, a phenotype believed to prevail at tumor initiation, whereas established tumors are dominated by Th2/M2-like macrophages^57^. These putative mechanisms were highlighted in our pathway analysis, with an enrichment of genes involved in INF-γ, PD-1 and IL-1 signaling among FEV_1_/FVC genetic instruments, and over-representation of pQTL targets in these pathways. Furthermore, a study of trans-thoracically implanted tumors in an emphysema mouse model demonstrates how this pulmonary phenotype results in impaired antitumor T cell responses at a critical point when nascent cancer cells evade detection and elimination by the immune system resulting in enhanced tumor growth^58^.

Other relevant pathways implicating pulmonary dysfunction in lung cancer development include lung tissue destruction via matrix degrading enzymes and increased genotoxic and apoptotic stress resulting from cigarette smoke in conjunction with macrophage- and neutrophil-derived ROS^15, 59^. This may explain our findings for FEV_1_ and squamous carcinoma, for which cigarette smoking is a particularly dominant risk factor. Genetic predisposition to impaired FEV_1_ may create a milieu that promotes malignant transformation and susceptibility to external carcinogens and tissue damage, rather than increasing the likelihood of cigarette smoking. In our analysis we attempted to isolate the former pathway from the latter by carefully instrumenting pulmonary phenotypes and confirming that they are not associated with behavioral aspects of nicotine dependence. However, residual confounding by smoking cannot be entirely precluded, given its high genetic and phenotypic correlation with FEV_1_.

The causal interpretation of our results critically depends on the validity of fundamental Mendelian randomization assumptions. We employed a range of estimation techniques with different underlying assumptions, as well as diagnostic tests, to interrogate the robustness of our results with respect to confounding, horizontal pleiotropy, and weak instrument bias. However, despite these efforts, residual confounding by related phenotypes, such as smoking, or subtle effects of population structure cannot be ruled out. In evaluating the contribution of our findings, several limitations should be acknowledged. Our approach to outlier removal based on Cochran’s Q statistic with modified second order weights may have been overly stringent; however, manually pruning based on such a large set of genetic instruments may not be feasible and may introduce additional bias, thus we feel this systematic conservative approach is justified. Furthermore, outlier removal did not have an adverse impact on instrument strength and precision of the MR analysis.

In addition to pleiotropy, selection bias may also undermine the validity of a Mendelian Randomization study, particularly in the form of collider bias, if selection is a function of the exposure or outcome. In the context of the UKB, low participation (5.5%) may have resulted in an unrepresentative study population^60,61^. Although enrolment in the cohort was not explicitly contingent on cancer status or pulmonary function, it is likely that individuals who did not complete a spirometry assessment were more likely to be smokers and have poor lung function. Simulations by Gkatzionis & Burgess^61^ demonstrate that when the effect of a risk factor on selection is mild to moderate (odds of selection: 0.82 to 0.61), the type I error rate remains reasonable at 5.0-6.6%. The direction of the resulting bias depends on the direction and strength of the exposure (lung function) – confounder (smoking) relationship. In the context of our study, the causal effect may be underestimated since the confounder and exposure are both likely to increase non-participation or result in missing spirometry data.

Another limitation is that we did not assess the relationship between the velocity of lung function decline and lung cancer risk, which may also prove to be a risk factor and capture a different dimension of pulmonary dysfunction. Furthermore, since our study includes the largest GWAS of lung cancer cases in never smokers, this precludes a well-powered replication study in an independent European ancestry population. In addition, dichotomous stratification by smoking status does not permit an evaluation of the relationship between pulmonary impairment and lung cancer risk across more granular levels of smoking. Lastly, in our efforts to present the most comprehensive assessment of pulmonary function impairment and lung cancer risk, a number of analyses were conducted, and it may be possible that some inconsistently observed associations were due to chance.

Despite these limitations, important strengths of this work include the large sample size for instrument development and causal hypothesis testing. Our Mendelian randomization approach leveraged a large number of genetic instruments, including variants specifically associated with lung function in never smokers, while balancing the concerns related to genetic confounding and pleiotropy. By triangulating evidence from gene expression and plasma protein levels, we also provide a more enriched interpretation of the genetic effects of pulmonary function loci on lung cancer risk, which implicate immune-mediated pathways. Despite the small individual SNP effect sizes, combining multiple instruments revealed meaningful increases in lung cancer risk. A genetically predicted 10% reduction in FEV_1_/FVC confers an approximately 55% increased risk of lung cancer in never smokers, and a similar magnitude of effect was observed for FEV_1_ and squamous carcinoma. However, effects of FEV_1_/FVC on adenocarcinoma were more modest (16-23% increase). Taken together, these findings provide more robust etiological insight than previous studies that relied on using observed lung function phenotypes directly as putatively casual factors.

As our understanding of the shared genetic and molecular pathways between lung cancer and pulmonary disease continues to evolve, identification of new susceptibility loci for pulmonary function and lung cancer risk may have important implications for future precision prevention and screening endeavors. Multiple genetic determinants of lung function are in pathways that contain druggable targets, based on our pQTL findings and previous reports^18^, which may open new avenues for chemoprevention or targeted therapies for lung cancers with an obstructive pulmonary etiology. In addition, with accumulating evidence supporting the effectiveness of low-dose computed tomography for lung cancer^62, 63^, impairment in FEV_1_ and FEV_1_/FVC and their genetic determinants may provide additional information for refining risk-stratification and screening eligibility criteria.

## METHODS

### Study Populations

The UK Biobank (UKB) is a population-based prospective cohort of over 500,000 individuals aged 40-69 years at enrollment in 2006-2010 who completed extensive questionnaires on health-related factors, physical assessments, and provided blood samples^64^. Participants were genotyped on the UK Biobank Affymetrix Axiom array (89%) or the UK BiLEVE array (11%)^64^. Genotype imputation was performed using the Haplotype Reference Consortium data as the main reference panel as well as using the merged UK10K and 1000 Genomes (1000G) phase 3 reference panels^64^. Our analyses were restricted to individuals of predominantly European ancestry based on self-report and after excluding samples with either of the first two genetic ancestry principal components (PCs) outside of 5 standard deviations (SD) of the population mean. Samples with discordant self-reported and genetic sex were removed. Using a subset of genotyped autosomal variants with minor allele frequency (MAF)≥0.01 and call rate ≥97%, we filtered samples with call rates <97% or heterozygosity >5 standard deviations (SD) from the mean. First-degree relatives were identified using KING^65^ and one sample from each pair was excluded, leaving at total of 413,810 individuals available for analysis.

We further excluded 36,461 individuals without spirometry data, 207 individuals who only completed one blow (n=207), for whom reproducibility could not be assessed (Supplementary Figure 1). For the remaining subjects, we examined the difference between the maximum value per individual (referred to as the best measure) and all other blows. Values differing by more than 0.15L were considered non-reproducible, based on standard spirometry guidelines^66^, and were excluded. Our analyses thus included 372,750 and 370,638 individuals for of FEV_1_ and FVC, respectively. The best per individual measure among the reproducible blows was used to derive FEV_1_/FVC, resulting in 368,817 individuals. FEV_1_ and FVC values were then converted to standardized Z-scores with a mean of 0 and standard deviation (SD) of 1.

The OncoArray Lung Cancer study has been previously described^16^. Briefly, this dataset consists of genome-wide summary statistics based on 29,266 lung cancer cases (11,273 adenocarcinoma, 7426 squamous carcinoma) and 56,450 controls of predominantly European ancestry (≥80%) assembled from studies part of the International Lung Cancer Consortium. Summary statistics from the lung cancer GWAS were adjusted for appropriate covariates, including genetic ancestry PCs, and showed no signs of genomic inflation for lung cancer overall (λ_GC_=1.0035) or for any subtypes, including adenocarcinoma (λ_GC_=1.0050), squamous carcinoma (λ_GC_=1.0051), and lung cancer in never smokers (λ_GC_=1.0060).

Informed consent was obtained from study participants in the UK Biobank and studies contributing data to the OncoArray Lung Cancer collaboration. UK Biobank received ethics approval from the Research Ethics Committee (REC reference: 11/NW/0382). Approval for OncoArray studies was obtained from each of the participating institutional research ethics review boards.

### Genome-Wide Association Analysis

Genome-wide association analyses of pulmonary function phenotypes in the UK Biobank cohort were conducted using PLINK 2.0 (October 2017 version). We excluded variants out of with Hardy-Weinberg equilibrium at p<1×10^−5^ in cancer-free individuals, call rate <95% (alternate allele dosage required to be within 0.1 of the nearest hard call to be non-missing), imputation quality INFO<0.30, and MAF<0.005. To minimize potential for reverse causation, prevalent lung cancer cases, defined as diagnoses occurring up to 5 years before cohort entry and incident cases occurring within 2 years of enrollment, were excluded (n=738). Linear regression models for pulmonary function phenotypes (standardized Z-scores for FEV_1_ and FVC; untransformed FEV_1_/FVC ratio bounded by 0 and 1) were adjusted for age, age^2^, sex, genotyping array and 15 PCs to permit an assessment of heritability (*h*_g_) and genetic correlation (*r*_g_) with height, smoking (status and pack-years), and anthropometric traits.

### Heritability and Genetic Correlation

LD Score regression^17^ was used to estimate *h*_g_ for each lung phenotype and *r*_g_ with lung cancer and other traits. To better capture LD patterns present in the UKB data, we generated LD scores for all variants that passed QC with MAF>0.0001 using a random sample of 10,000 UKB participants. UKB LD scores were used to estimate *h*_g_ for each lung phenotype and *r*_g_ with other non-cancer traits. Genetic correlation with lung cancer was estimated using publicly available LD scores based on the 1000G phase 3 reference population (n=1,095,408 variants).

To assess the importance of specific functional annotations in SNP-heritability, we partitioned trait-specific heritability using stratified-LDSC^67^. The analysis was performed using 86 annotations (baseline-LD model v2.1), which incorporated MAF-adjustment and other LD-related annotations, such as predicted allele age and recombination rate^20, 22^. The MHC region was excluded from partitioned heritability analyses. Enrichment was considered statistically significant if p<8.5×10^−4^, which reflects Bonferroni correction for 59 annotations (functional categories with and without a 500 bp window around it were considered as the same annotation).

### Development of Genetic Instruments for Pulmonary Function

For the purpose of instrument development, a two-stage genome-wide analysis was employed, with a randomly sampled 70% of the cohort used for discovery and the remaining 30% reserved for replication. In addition to age, age^2^, sex, genotyping array and 15 PC’s, models were adjusted for covariates that explain a substantial proportion of variation in pulmonary phenotypes, such as smoking and height, in order to decrease the residual variance and help isolate the relevant genetic signals. Specifically, we adjusted for height, height^2^, and cigarette pack-year categories (0, corresponding to never-smokers, >0-10, >10-20, >20-30, >30-40, and >40). Other covariates, such as UKB assessment center (Field 54), use of an inhaler prior to spirometry (Field 3090), and blow acceptability (Field 3061) were considered. However, these covariates did not explain a substantial proportion of phenotype variation and had low variable importance metrics (lmg<0.01), and thus were not included in our final models. Instruments were selected from independent associated variants (LD *r*^2^<0.05 in a clumping window of 10,000 kb) with *P*<5×10^−8^ in the discovery stage and *P*<0.05 and consistent direction of effect in the replication stage. Since the primary goal of our GWAS was to develop a comprehensive set of genetic instruments we applied a less stringent replication threshold in anticipation of subsequent filtering based on potential violation of Mendelian randomization assumptions.

### Mendelian Randomization

Mendelian randomization (MR) analyses were carried out to investigate the potential causal relationship between impaired pulmonary function and lung cancer risk. Genetic instruments excluded multi-allelic and non-inferable palindromic variants with intermediate allele frequencies (MAF>0.42). Odds ratios (OR) and corresponding 95% confidence intervals were obtained using the maximum likelihood and inverse variance weighted multiplicative random-effects (IVW-RE) estimators^28, 29^. Effects for FEV_1_ and FVC were estimated for a genetically predicted 1-SD decrease in the standardized Z-score. For FEV_1_/FVC, we modeled cancer risk corresponding to a 10% decrease in the ratio. Sensitivity analyses included the weighted median (WM) estimator^30^, which provides unbiased estimates when up to 50% of the weights are from invalid instruments, and MR RAPS (Robust Adjusted Profile Score), which incorporates random effect and robust loss functions to limit the influence of potentially pleiotropic instruments. MR RAPS assumes balanced (mean 0) horizontal pleiotropy. In contrast to IVW-RE, MR RAPS models idiosyncratic and systematic pleiotropy effects as additive, rather than multiplicative^31^. Using MR estimation techniques with different underlying statistical models allows for a more comprehensive assessment of the robustness of our results with respect to violations of MR assumptions. We also applied the following diagnostic tests: i) significant (p<0.05) deviation of the MR Egger intercept (β_0 *Egger*_) from 0, as a test for directional pleiotropy^68^; ii) *I*^2^_GX_ statistic <0.90 indicative of regression dilution bias and inflation in the MR Egger pleiotropy test due to violation of the no measurement error (NOME) assumption^68^; iii) Cochran’s Q-statistic with modified second order weights to asses heterogeneity (p-value<0.05) indicative of (balanced) horizontal pleiotropy^69^.

All statistical analyses were conducted using R (version 3.6.1). Mendelian randomization analyses were conducted using the TwoSampleMR R package (version 0.4.23).

### Functional Characterization of Lung Function Instruments

In order to characterize functional pathways that are represented by the genetic instruments for FEV_1_ and FEV_1_/FVC, we examined effects on gene expression in lung tissues from 409 subjects from the Laval eQTL study^35^. Lung function instruments with significant (Bonferroni p-value<0.05) eQTL effects were used as instruments to estimate the effect of the gene expression on lung cancer risk. For genes with multiple eQTLs, independent variants (LD *r*^2^<0.05) were used to obtain IVW estimates of the predicted effects of increased gene expression on lung cancer risk. For genes with a single eQTL, OR estimates were obtained using the Wald method. Next, we examined data from the genetic atlas of the human plasma proteome^36^, queried using PhenoScanner^70^, to assess whether any of the genetic instruments for FEV_1_ and FEV_1_/FVC had significant (p<5×10^−8^) effects on intracellular protein levels. Lastly, we summarized the pathways represented by the genes where the lung function instruments were localized using pathway enrichment analysis via the Reactome database and ImmuneSigDB (collection C7 from MSigDB).

## DATA AVAILABILITY

The datasets generated during and/or analyzed during the current study are available from the authors on request. Genotype data for the Oncoarray Consortium Lung Cancer studies have been deposited in the database of Genotypes and Phenotypes (dbGaP) under accession: phs001273.v2.p2 [https://www.ncbi.nlm.nih.gov/projects/gap/cgi-bin/study.cgi?study_id=phs001273.v3.p2]. Readers interested in obtaining a copy of the lung cancer GWAS summary statistics can do so by completing the proposal request form at http://oncoarray.dartmouth.edu/. The UK Biobank in an open access resource, available at https://www.ukbiobank.ac.uk/researchers/. This research was conducted with approved access to UK Biobank data under applications number 14105 and 23261. All data supporting the findings of this study are available within the article and its supplementary information files, and from the corresponding authors upon reasonable request. A reporting summary for this article is available as a Supplementary Information file.

## URLs

PLINK 2.0: https://www.cog-genomics.org/plink/2.0/

LDSC (version 1.0.0) from: https://github.com/bulik/ldsc/

LDSC functional annotations available from: https://data.broadinstitute.org/alkesgroup/LDSCORE/1000G_Phase3_EUR_baselineLD_v2.1_ldscores.tgz

R package for Circos plots (version 0.4.7): https://github.com/jokergoo/circlize

R package for Mendelian Randomization (version 0.4.23): https://github.com/MRCIEU/TwoSampleMR R package for PhenoScanner (version 1.0): https://github.com/phenoscanner/phenoscanner

R packages for pathway analysis: https://bioconductor.org/packages/release/bioc/html/ReactomePA.html and https://bioconductor.org/packages/release/bioc/html/clusterProfiler.html

ImmuneSigDB (C7): http://software.broadinstitute.org/gsea/msigdb/collections.jsp

## ACKNOWEDGEMENTS

Disclaimer: Where authors are identified as personnel of the International Agency for Research on Cancer / World Health Organization, the authors alone are responsible for the views expressed in this article and they do not necessarily represent the decisions, policy or views of the International Agency for Research on Cancer / World Health Organization.

This research was supported by funding from the National Institutes of Health (US NCI R25T CA112355 and R01 CA201358; PI: Witte) and the Canadian Institute for Health Research (Foundation grant FDN 167273, PI: Hung; Canada Research Chair, PI: Hung).

The OncoArray project was supported by NIH U19 CA203654 (MPI: Hung, Amos, Brennan, Lin). The Boston Lung Cancer Study was funded by NIH (NCI) U01CA209414 (PI: Christiani).

The lung eQTL study at Laval University was supported by the Fondation de l’Institut universitaire de cardiologie et de pneumologie de Québec and the Canadian Institutes of Health Research (MOP - 123369). Y.B. holds a Canada Research Chair in Genomics of Heart and Lung Diseases.

The EAGLE study was supported by the Intramural Research Program, Division of Cancer Epidemiology and Genetics, National Cancer Institute, NIH, DHHS.

The Multiethnic Cohort Study is supported by National Institutes of Health (CA164973).

The CARET study was supported by the National Institutes of Health / National Cancer Institute: UM1 CA167462 (PI: Goodman), U01 CA6367307 (PIs: Omen, Goodman); R01 CA111703 (PI: Chen), 5R01 CA151989-01A1 (PI: Doherty) and U01 CA167462 (PI: Chen).

Matthew B. Schabath was supported in part by a Cancer Center Support Grant (P30 CA076292) and by NIH P50 CA119997.

Richard M. Martin is supported by a CRUK programme grant, the Integrative Cancer Epidemiology Programme (C18281/A19169), and by the National Institute for Health Research (NIHR) Bristol Biomedical Research Centre based at University Hospitals Bristol NHS Foundation Trust and the University of Bristol. The views expressed in this publication are those of the authors and not necessarily those of the NHS, the National Institute for Health Research or the Department of Health.

Dr Haycock is supported by CRUK Population Research Postdoctoral Fellowship C52724/A20138.

Michael P. Davies is supported by the Roy Castle Lung Cancer Foundation UK.

## AUTHOR CONTRIBUTIONS

Study conception: L.K., R.J.H, M.J., P.B.; Statistical analysis: L.K.; L.K drafted the manuscript with input from R.J.H. and J.S.W; Project coordination: R.J.H, P.B., M.J., J.S.W.; UK Biobank genotype and sample quality control: S.R.R., R.E.G; Lung tissue eQTL analysis: Y. Bossé, V.M.; Development of analytic strategy: L.K., S.R.R., R.E.G., R.J.H., J.S.W., M.J., R.M.M., C.R., G.D.S., P.C.H.; Data acquisition and development of lung cancer epidemiological studies: R.J.H., C.I.A., P.B., M.J., J.S.W., N.E.C., M.T.L., D.C.C., P.V., G.L., G.S., D.Z., S.S.S., D.A., M.C.A., A.T., G.R., C.C., G.E.G, J.A.D., H.B., J.K.F, M.P.D., M.D.T., L.A.K., S.E.B., A.H., S.Z., S.L., L.L.M., I.C., M.B.S., E.J.D., A.S.A., J.M., P.L., S.A., J.D.M., N.C.E., M.T.W., Y.B., M.O., Y. Bossé. All authors contributed to the interpretation of the results and provided critical feedback on the manuscript.

