## Supplementary Tables and Figures for "Immune-mediated genetic pathways resulting in pulmonary function impairment increase lung cancer susceptibility"

**Supplementary Table 1:** Functional partitioning of array-based heritability for $FEV_1$, based on genome-wide summary statistics from the UK Biobank cohort and UK Biobank-specific LD scores.

| Category | Pr(SNPs) | Pr($h_g$) | SE | Enrichment | SE | Enrichment P |
|---|---|---|---|---|---|---|
| H3K4me1 Trynka (500bp window) | 0.606 | 0.892 | 0.024 | 1.473 | 0.039 | 1.86E-24 |
| SuperEnhancer Hnisz (500bp window) | 0.170 | 0.339 | 0.014 | 1.989 | 0.084 | 3.35E-24 |
| CpG Content 50kb | 0.010 | 0.012 | 0.0002 | 1.201 | 0.017 | 7.64E-24 |
| H3K27ac Hnisz (500bp window) | 0.420 | 0.676 | 0.023 | 1.609 | 0.054 | 8.05E-23 |
| H3K27ac Hnisz | 0.389 | 0.666 | 0.023 | 1.712 | 0.060 | 1.02E-22 |
| Background Selection Statistic | 0.178 | 0.242 | 0.006 | 1.364 | 0.034 | 1.91E-20 |
| SuperEnhancer Hnisz | 0.167 | 0.333 | 0.017 | 1.992 | 0.099 | 1.70E-19 |
| Conserved Primate phastCons (500bp window) | 0.176 | 0.568 | 0.041 | 3.230 | 0.232 | 3.12E-18 |
| Conserved LindbladToh (500bp window) | 0.330 | 0.617 | 0.031 | 1.866 | 0.095 | 1.14E-16 |
| Conserved Primate phastCons | 0.019 | 0.251 | 0.023 | 13.022 | 1.207 | 1.58E-16 |
| H3K9ac Trynka | 0.125 | 0.488 | 0.041 | 3.889 | 0.331 | 3.41E-16 |
| Conserved Mammal phastCons (500bp window) | 0.339 | 0.674 | 0.041 | 1.986 | 0.121 | 8.07E-14 |
| H3K27ac PGC2 (500bp window) | 0.335 | 0.567 | 0.032 | 1.690 | 0.095 | 5.03E-13 |
| H3K9ac Trynka (500bp window) | 0.230 | 0.518 | 0.040 | 2.256 | 0.173 | 5.44E-13 |
| Intron UCSC (500bp window) | 0.397 | 0.514 | 0.015 | 1.296 | 0.037 | 2.85E-12 |
| DGF ENCODE (500bp window) | 0.538 | 0.858 | 0.043 | 1.593 | 0.081 | 1.78E-11 |
| Bivariate Flanking TSS/Enhancer | 0.014 | 0.111 | 0.015 | 8.183 | 1.083 | 1.33E-10 |
| Conserved LindbladToh | 0.026 | 0.306 | 0.040 | 11.923 | 1.575 | 1.98E-10 |
| H3K4me1 Trynka | 0.424 | 0.760 | 0.053 | 1.793 | 0.125 | 5.71E-10 |
| Conserved Vertebrate phastCons | 0.029 | 0.261 | 0.035 | 8.861 | 1.181 | 7.37E-10 |
| H3K4me3 Trynka (500bp window) | 0.255 | 0.524 | 0.043 | 2.053 | 0.169 | 1.16E-09 |
| Enhancer Hoffman (500bp window) | 0.090 | 0.238 | 0.025 | 2.644 | 0.276 | 2.57E-09 |
| Conserved Mammal phastCons | 0.021 | 0.234 | 0.034 | 10.893 | 1.596 | 4.66E-09 |
| H3K4me3 Trynka | 0.133 | 0.341 | 0.035 | 2.561 | 0.260 | 5.42E-09 |
| H3K27ac PGC2 | 0.269 | 0.520 | 0.043 | 1.935 | 0.160 | 1.32E-08 |
| TSS Hoffman (500bp window) | 0.034 | 0.125 | 0.016 | 3.629 | 0.458 | 1.58E-08 |
| DHS Trynka (500bp window) | 0.496 | 0.754 | 0.045 | 1.521 | 0.091 | 5.49E-08 |
| Conserved Vertebrate phastCons (500bp window) | 0.407 | 0.655 | 0.044 | 1.610 | 0.107 | 6.69E-08 |
| TFBS ENCODE (500bp window) | 0.341 | 0.587 | 0.046 | 1.720 | 0.136 | 2.97E-07 |
| DGF ENCODE | 0.136 | 0.449 | 0.057 | 3.302 | 0.422 | 3.70E-07 |
| Fetal DHS Trynka (500bp window) | 0.283 | 0.502 | 0.046 | 1.773 | 0.162 | 5.83E-06 |
| BLUEPRINT DNA methylation MaxCPP | 0.032 | 0.100 | 0.014 | 3.158 | 0.444 | 6.00E-06 |
| Bivariate Flanking TSS/Enhancer (500bp window) | 0.031 | 0.118 | 0.019 | 3.797 | 0.625 | 1.03E-05 |
| TSS Hoffman | 0.018 | 0.094 | 0.017 | 5.287 | 0.967 | 1.48E-05 |
| Coding UCSC | 0.014 | 0.098 | 0.018 | 6.873 | 1.286 | 1.63E-05 |
| Intron UCSC | 0.387 | 0.459 | 0.017 | 1.185 | 0.043 | 2.19E-05 |
| DHS Trynka | 0.166 | 0.438 | 0.060 | 2.636 | 0.363 | 2.29E-05 |

| | | | | | | |
|---|---|---|---|---|---|---|
| DHS peaks Trynka | 0.111 | 0.344 | 0.057 | 3.114 | 0.519 | 9.96E-05 |
| Non-synonymous | 0.003 | 0.037 | 0.008 | 13.481 | 3.069 | 1.07E-04 |
| TFBS ENCODE | 0.131 | 0.331 | 0.051 | 2.524 | 0.392 | 1.10E-04 |
| Enhancer Hoffman | 0.042 | 0.153 | 0.030 | 3.649 | 0.707 | 2.36E-04 |
| UTR 3 UCSC | 0.011 | 0.078 | 0.018 | 7.006 | 1.613 | 2.79E-04 |
| H3K4me1 peaks Trynka | 0.170 | 0.374 | 0.056 | 2.204 | 0.329 | 3.01E-04 |
| UTR 3 UCSC (500bp window) | 0.026 | 0.085 | 0.017 | 3.229 | 0.626 | 4.75E-04 |
| Coding UCSC (500bp window) | 0.064 | 0.123 | 0.017 | 1.934 | 0.261 | 5.29E-04 |
| Fetal DHS Trynka | 0.084 | 0.256 | 0.049 | 3.051 | 0.582 | 6.17E-04 |
| H3K4me3 peaks Trynka | 0.042 | 0.144 | 0.033 | 3.468 | 0.800 | 1.99E-03 |
| GERP.RSsup4 | 0.008 | 0.080 | 0.024 | 9.874 | 2.933 | 3.07E-03 |
| H3K9ac peaks Trynka | 0.038 | 0.147 | 0.037 | 3.824 | 0.955 | 3.34E-03 |
| GTEx eQTL MaxCPP | 0.010 | 0.040 | 0.010 | 3.889 | 1.010 | 5.75E-03 |
| BLUEPRINT H3K27acQTL MaxCPP | 0.017 | 0.048 | 0.013 | 2.898 | 0.815 | 0.022 |
| BLUEPRINT H3K4me1QTL MaxCPP | 0.013 | 0.035 | 0.010 | 2.645 | 0.768 | 0.032 |
| UTR 5 UCSC | 0.005 | 0.027 | 0.010 | 4.881 | 1.761 | 0.033 |
| Promoter UCSC (500bp window) | 0.057 | 0.084 | 0.017 | 1.482 | 0.304 | 0.113 |
| Promoter UCSC | 0.046 | 0.074 | 0.019 | 1.607 | 0.404 | 0.132 |
| Enhancer Andersson | 0.004 | 0.019 | 0.010 | 4.472 | 2.365 | 0.144 |
| Promoter Flanking Hoffman (500bp window) | 0.033 | 0.060 | 0.019 | 1.812 | 0.583 | 0.163 |
| CTCF Hoffman (500bp window) | 0.071 | 0.099 | 0.026 | 1.392 | 0.368 | 0.289 |
| Enhancer Andersson (500bp window) | 0.019 | 0.030 | 0.012 | 1.566 | 0.617 | 0.358 |
| Transcr Hoffman | 0.346 | 0.389 | 0.047 | 1.123 | 0.136 | 0.369 |
| Synonymous | 0.003 | 0.012 | 0.011 | 3.791 | 3.561 | 0.433 |
| UTR 5 UCSC (500bp window) | 0.027 | 0.034 | 0.014 | 1.275 | 0.515 | 0.594 |
| Weak Enhancer Hoffman | 0.021 | 0.018 | 0.020 | 0.837 | 0.975 | 0.867 |
| CTCF Hoffman | 0.024 | 0.026 | 0.022 | 1.110 | 0.928 | 0.906 |
| Weak Enhancer Hoffman (500bp window) | 0.089 | 0.089 | 0.026 | 1.001 | 0.296 | 0.998 |

**Supplementary Table 2**: Functional partitioning of array-based heritability for $FEV_1$/FVC, based on genome-wide summary statistics from the UK Biobank cohort and UK Biobank-specific LD scores.

| Category | Pr(SNPs) | Pr($h_g$) | SE | Enrichment | SE | Enrichment P |
|---|---|---|---|---|---|---|
| H3K27ac Hnisz | 0.389 | 0.800 | 0.030 | 2.055 | 0.077 | 4.18E-23 |
| SuperEnhancer Hnisz | 0.167 | 0.386 | 0.020 | 2.309 | 0.119 | 9.57E-22 |
| SuperEnhancer Hnisz (500bp window) | 0.170 | 0.397 | 0.019 | 2.330 | 0.111 | 1.31E-21 |
| H3K27ac Hnisz (500bp window) | 0.420 | 0.768 | 0.031 | 1.826 | 0.075 | 4.48E-21 |
| H3K4me1 Trynka (500bp window) | 0.606 | 0.922 | 0.032 | 1.522 | 0.052 | 5.38E-19 |
| CpG Content 50kb | 0.010 | 0.012 | 0.0002 | 1.187 | 0.020 | 3.84E-17 |
| Background Selection Statistic | 0.178 | 0.226 | 0.006 | 1.272 | 0.035 | 1.51E-15 |
| H3K27ac PGC2 (500bp window) | 0.335 | 0.612 | 0.035 | 1.825 | 0.106 | 9.04E-15 |
| H3K9ac Trynka | 0.125 | 0.485 | 0.047 | 3.871 | 0.378 | 5.98E-13 |
| H3K9ac Trynka (500bp window) | 0.230 | 0.607 | 0.051 | 2.639 | 0.221 | 2.00E-12 |
| Conserved Primate phastCons (500bp window) | 0.176 | 0.547 | 0.050 | 3.109 | 0.282 | 2.15E-12 |
| H3K4me3 Trynka (500bp window) | 0.255 | 0.616 | 0.050 | 2.413 | 0.195 | 3.83E-12 |
| Fetal DHS Trynka (500bp window) | 0.283 | 0.655 | 0.050 | 2.311 | 0.178 | 7.85E-11 |
| Enhancer Hoffman (500bp window) | 0.090 | 0.279 | 0.030 | 3.107 | 0.329 | 4.02E-10 |
| DGF ENCODE (500bp window) | 0.538 | 0.862 | 0.049 | 1.601 | 0.092 | 7.56E-10 |
| Conserved Mammal phastCons (500bp window) | 0.339 | 0.702 | 0.058 | 2.070 | 0.171 | 1.74E-09 |
| H3K4me1 Trynka | 0.424 | 0.800 | 0.059 | 1.888 | 0.139 | 4.28E-09 |
| H3K4me3 Trynka | 0.133 | 0.364 | 0.041 | 2.739 | 0.306 | 1.17E-08 |
| Fetal DHS Trynka | 0.084 | 0.420 | 0.059 | 5.010 | 0.698 | 1.33E-07 |
| Bivariate Flanking TSS/Enhancer | 0.014 | 0.131 | 0.022 | 9.685 | 1.638 | 2.47E-07 |
| Conserved Vertebrate phastCons (500bp window) | 0.407 | 0.686 | 0.056 | 1.686 | 0.136 | 5.00E-07 |
| DHS Trynka (500bp window) | 0.496 | 0.777 | 0.056 | 1.566 | 0.114 | 7.71E-07 |
| Conserved LindbladToh | 0.026 | 0.212 | 0.038 | 8.239 | 1.473 | 9.52E-07 |
| Enhancer Hoffman | 0.042 | 0.211 | 0.033 | 5.014 | 0.776 | 1.02E-06 |
| Intron UCSC (500bp window) | 0.397 | 0.500 | 0.020 | 1.259 | 0.051 | 1.95E-06 |
| Conserved LindbladToh (500bp window) | 0.330 | 0.583 | 0.056 | 1.763 | 0.169 | 3.49E-06 |
| H3K27ac PGC2 | 0.269 | 0.512 | 0.053 | 1.905 | 0.199 | 4.17E-06 |
| Conserved Primate phastCons | 0.019 | 0.165 | 0.031 | 8.548 | 1.610 | 4.91E-06 |
| Bivariate Flanking TSS/Enhancer (500bp window) | 0.031 | 0.126 | 0.022 | 4.050 | 0.691 | 1.17E-05 |
| Conserved Mammal phastCons | 0.021 | 0.151 | 0.032 | 7.057 | 1.480 | 2.48E-05 |
| H3K4me1 peaks Trynka | 0.170 | 0.473 | 0.071 | 2.785 | 0.416 | 2.62E-05 |
| DHS peaks Trynka | 0.111 | 0.532 | 0.097 | 4.809 | 0.875 | 3.06E-05 |
| DHS Trynka | 0.166 | 0.584 | 0.095 | 3.515 | 0.574 | 3.35E-05 |
| DGF ENCODE | 0.136 | 0.462 | 0.079 | 3.398 | 0.582 | 9.07E-05 |
| Conserved Vertebrate phastCons | 0.029 | 0.148 | 0.033 | 5.028 | 1.108 | 2.27E-04 |
| TFBS ENCODE (500bp window) | 0.341 | 0.525 | 0.050 | 1.539 | 0.146 | 2.51E-04 |
| GTEx eQTL MaxCPP | 0.010 | 0.054 | 0.013 | 5.239 | 1.216 | 3.94E-04 |

| | | | | | | |
|---|---|---|---|---|---|---|
| BLUEPRINT DNA methylation MaxCPP | 0.032 | 0.102 | 0.019 | 3.206 | 0.585 | 4.10E-04 |
| TSS Hoffman (500bp window) | 0.034 | 0.110 | 0.022 | 3.213 | 0.630 | 5.89E-04 |
| H3K9ac peaks Trynka | 0.038 | 0.199 | 0.049 | 5.168 | 1.279 | 8.69E-04 |
| H3K4me3 peaks Trynka | 0.042 | 0.170 | 0.042 | 4.093 | 1.011 | 1.82E-03 |
| Intron UCSC | 0.387 | 0.465 | 0.027 | 1.200 | 0.069 | 4.87E-03 |
| Coding UCSC (500bp window) | 0.064 | 0.147 | 0.030 | 2.301 | 0.475 | 6.36E-03 |
| TFBS ENCODE | 0.131 | 0.311 | 0.067 | 2.373 | 0.514 | 7.18E-03 |
| UTR 3 UCSC | 0.011 | 0.048 | 0.014 | 4.325 | 1.226 | 7.51E-03 |
| TSS Hoffman | 0.018 | 0.079 | 0.024 | 4.407 | 1.326 | 9.21E-03 |
| Coding UCSC | 0.014 | 0.059 | 0.019 | 4.125 | 1.332 | 0.018 |
| BLUEPRINT H3K4me1QTL MaxCPP | 0.013 | 0.037 | 0.010 | 2.759 | 0.743 | 0.019 |
| GERP.RSsup4 | 0.008 | 0.061 | 0.025 | 7.448 | 3.036 | 0.031 |
| Enhancer Andersson | 0.004 | 0.031 | 0.013 | 7.128 | 2.967 | 0.043 |
| UTR 3 UCSC (500bp window) | 0.026 | 0.066 | 0.020 | 2.482 | 0.766 | 0.052 |
| BLUEPRINT H3K27acQTL MaxCPP | 0.017 | 0.045 | 0.015 | 2.735 | 0.888 | 0.053 |
| Non-synonymous | 0.003 | 0.016 | 0.008 | 5.847 | 3.040 | 0.117 |
| UTR 5 UCSC (500bp window) | 0.027 | 0.045 | 0.017 | 1.669 | 0.619 | 0.274 |
| Transcr Hoffman | 0.346 | 0.408 | 0.058 | 1.179 | 0.168 | 0.293 |
| WeakEnhancer Hoffman (500bp window) | 0.089 | 0.120 | 0.036 | 1.352 | 0.403 | 0.379 |
| WeakEnhancer Hoffman | 0.021 | 0.041 | 0.024 | 1.960 | 1.145 | 0.398 |
| Transcr Hoffman (500bp window) | 0.762 | 0.729 | 0.041 | 0.956 | 0.054 | 0.419 |
| CTCF Hoffman | 0.024 | 0.005 | 0.028 | 0.206 | 1.192 | 0.504 |
| PromoterFlanking Hoffman (500bp window) | 0.033 | 0.049 | 0.024 | 1.467 | 0.718 | 0.514 |
| Promoter UCSC (500bp window) | 0.057 | 0.068 | 0.022 | 1.192 | 0.381 | 0.614 |
| CTCF Hoffman (500bp window) | 0.071 | 0.058 | 0.031 | 0.812 | 0.443 | 0.670 |
| Enhancer Andersson (500bp window) | 0.019 | 0.026 | 0.019 | 1.342 | 1.006 | 0.733 |
| UTR 5 UCSC | 0.005 | 0.009 | 0.014 | 1.558 | 2.638 | 0.832 |
| Synonymous | 0.003 | 0.001 | 0.014 | 0.457 | 4.364 | 0.901 |
| Promoter UCSC | 0.046 | 0.048 | 0.026 | 1.035 | 0.550 | 0.950 |

**Supplementary Table 3:** Functional partitioning of array-based heritability for FVC, based on genome-wide summary statistics from the UK Biobank cohort and UK Biobank-specific LD scores.

| Category | Pr(SNPs) | Pr(h2) | SE | Enrichment | SE | Enrichment P |
|---|---|---|---|---|---|---|
| CpG Content 50kb | 0.010 | 0.012 | 0.0002 | 1.230 | 0.019 | 2.34E-23 |
| Background Selection Statistic | 0.178 | 0.251 | 0.006 | 1.411 | 0.036 | 6.87E-23 |
| SuperEnhancer Hnisz (500bp window) | 0.170 | 0.347 | 0.016 | 2.037 | 0.095 | 5.12E-20 |
| H3K4me1 Trynka (500bp window) | 0.606 | 0.893 | 0.027 | 1.474 | 0.045 | 9.09E-20 |
| Conserved Primate phastCons (500bp window) | 0.176 | 0.585 | 0.042 | 3.325 | 0.236 | 2.08E-18 |
| H3K27ac Hnisz | 0.389 | 0.652 | 0.027 | 1.675 | 0.069 | 2.80E-18 |
| H3K27ac Hnisz (500bp window) | 0.420 | 0.666 | 0.026 | 1.583 | 0.062 | 9.25E-18 |
| SuperEnhancer Hnisz | 0.167 | 0.344 | 0.018 | 2.057 | 0.109 | 7.54E-17 |
| Conserved Primate phastCons | 0.019 | 0.253 | 0.025 | 13.125 | 1.290 | 1.91E-15 |
| H3K9ac Trynka (500bp window) | 0.230 | 0.575 | 0.040 | 2.503 | 0.175 | 2.37E-15 |
| Conserved LindbladToh (500bp window) | 0.330 | 0.648 | 0.036 | 1.961 | 0.110 | 6.60E-15 |
| H3K9ac Trynka | 0.125 | 0.481 | 0.046 | 3.839 | 0.367 | 6.48E-13 |
| Intron UCSC (500bp window) | 0.397 | 0.527 | 0.016 | 1.328 | 0.040 | 1.29E-12 |
| Conserved Mammal phastCons (500bp window) | 0.339 | 0.706 | 0.048 | 2.082 | 0.140 | 1.51E-12 |
| H3K27ac PGC2 (500bp window) | 0.335 | 0.585 | 0.036 | 1.746 | 0.107 | 3.34E-11 |
| DGF ENCODE (500bp window) | 0.538 | 0.874 | 0.046 | 1.623 | 0.085 | 3.53E-11 |
| Conserved LindbladToh | 0.026 | 0.297 | 0.039 | 11.556 | 1.517 | 1.51E-10 |
| H3K4me3 Trynka (500bp window) | 0.255 | 0.538 | 0.043 | 2.110 | 0.169 | 3.76E-10 |
| Conserved Mammal phastCons | 0.021 | 0.242 | 0.032 | 11.282 | 1.506 | 3.80E-10 |
| H3K4me1 Trynka | 0.424 | 0.769 | 0.052 | 1.816 | 0.123 | 5.46E-10 |
| Conserved Vertebrate phastCons | 0.029 | 0.254 | 0.034 | 8.630 | 1.148 | 9.43E-10 |
| H3K4me3 Trynka | 0.133 | 0.350 | 0.035 | 2.634 | 0.266 | 4.30E-09 |
| Enhancer Hoffman (500bp window) | 0.090 | 0.249 | 0.026 | 2.776 | 0.295 | 5.72E-09 |
| Bivariate Flanking TSS/Enhancer | 0.014 | 0.132 | 0.020 | 9.766 | 1.447 | 8.44E-09 |
| DGF ENCODE | 0.136 | 0.506 | 0.060 | 3.724 | 0.443 | 1.58E-08 |
| DHS Trynka (500bp window) | 0.496 | 0.778 | 0.047 | 1.568 | 0.094 | 2.63E-08 |
| TSS Hoffman (500bp window) | 0.034 | 0.137 | 0.019 | 4.002 | 0.557 | 1.51E-07 |
| Conserved Vertebrate phastCons (500bp window) | 0.407 | 0.681 | 0.050 | 1.674 | 0.124 | 1.67E-07 |
| H3K27ac PGC2 | 0.269 | 0.513 | 0.045 | 1.909 | 0.168 | 2.47E-07 |
| TSS Hoffman | 0.018 | 0.110 | 0.019 | 6.152 | 1.093 | 4.03E-06 |
| TFBS ENCODE (500bp window) | 0.341 | 0.590 | 0.052 | 1.728 | 0.153 | 4.54E-06 |
| TFBS ENCODE | 0.131 | 0.392 | 0.056 | 2.989 | 0.428 | 8.29E-06 |
| DHS Trynka | 0.166 | 0.466 | 0.064 | 2.805 | 0.386 | 1.14E-05 |
| Bivariate Flanking TSS/Enhancer (500bp window) | 0.031 | 0.132 | 0.023 | 4.223 | 0.729 | 1.51E-05 |
| DHS peaks Trynka | 0.111 | 0.374 | 0.061 | 3.386 | 0.554 | 3.68E-05 |
| BLUEPRINT DNA methylation MaxCPP | 0.032 | 0.101 | 0.016 | 3.169 | 0.508 | 4.61E-05 |
| Coding UCSC | 0.014 | 0.102 | 0.021 | 7.153 | 1.450 | 5.50E-05 |

| | | | | | | |
|---|---|---|---|---|---|---|
| Intron UCSC | 0.387 | 0.461 | 0.019 | 1.191 | 0.048 | 1.09E-04 |
| H3K4me1 peaks Trynka | 0.170 | 0.379 | 0.055 | 2.233 | 0.321 | 1.75E-04 |
| Coding UCSC (500bp window) | 0.064 | 0.143 | 0.021 | 2.241 | 0.323 | 2.21E-04 |
| FetalDHS Trynka (500bp window) | 0.283 | 0.482 | 0.053 | 1.702 | 0.185 | 2.52E-04 |
| Enhancer Hoffman | 0.042 | 0.160 | 0.034 | 3.822 | 0.804 | 6.03E-04 |
| UTR 3 UCSC | 0.011 | 0.082 | 0.020 | 7.327 | 1.823 | 7.48E-04 |
| Non-synonymous | 0.003 | 0.033 | 0.009 | 12.121 | 3.398 | 1.42E-03 |
| FetalDHS Trynka | 0.084 | 0.243 | 0.049 | 2.894 | 0.587 | 1.51E-03 |
| UTR 3 UCSC (500bp window) | 0.026 | 0.081 | 0.018 | 3.076 | 0.675 | 2.41E-03 |
| GERP.RSsup4 | 0.008 | 0.077 | 0.022 | 9.487 | 2.753 | 2.63E-03 |
| GTEx eQTL MaxCPP | 0.010 | 0.045 | 0.012 | 4.395 | 1.140 | 3.69E-03 |
| BLUEPRINT H3K27acQTL MaxCPP | 0.017 | 0.061 | 0.015 | 3.658 | 0.899 | 4.00E-03 |
| H3K4me3 peaks Trynka | 0.042 | 0.137 | 0.036 | 3.291 | 0.861 | 8.58E-03 |
| UTR 5 UCSC | 0.005 | 0.033 | 0.012 | 5.952 | 2.178 | 0.025 |
| BLUEPRINT H3K4me1QTL MaxCPP | 0.013 | 0.037 | 0.011 | 2.767 | 0.811 | 0.030 |
| H3K9ac peaks Trynka | 0.038 | 0.125 | 0.044 | 3.248 | 1.131 | 0.049 |
| PromoterFlanking Hoffman (500bp window) | 0.033 | 0.074 | 0.022 | 2.230 | 0.669 | 0.068 |
| Promoter UCSC (500bp window) | 0.057 | 0.088 | 0.019 | 1.547 | 0.331 | 0.099 |
| Promoter UCSC | 0.046 | 0.076 | 0.020 | 1.639 | 0.429 | 0.137 |
| Synonymous | 0.003 | 0.023 | 0.014 | 7.340 | 4.368 | 0.150 |
| Transcr Hoffman | 0.346 | 0.419 | 0.052 | 1.211 | 0.150 | 0.162 |
| Enhancer Andersson (500bp window) | 0.019 | 0.037 | 0.013 | 1.966 | 0.699 | 0.165 |
| UTR 5 UCSC (500bp window) | 0.027 | 0.046 | 0.014 | 1.727 | 0.536 | 0.179 |
| Enhancer Andersson | 0.004 | 0.019 | 0.011 | 4.303 | 2.560 | 0.198 |
| WeakEnhancer Hoffman | 0.021 | 0.037 | 0.024 | 1.762 | 1.127 | 0.498 |
| CTCF Hoffman | 0.024 | 0.028 | 0.026 | 1.173 | 1.071 | 0.872 |
| WeakEnhancer Hoffman (500bp window) | 0.089 | 0.093 | 0.028 | 1.050 | 0.311 | 0.872 |
| CTCF Hoffman (500bp window) | 0.071 | 0.073 | 0.027 | 1.026 | 0.379 | 0.946 |

**Supplementary Table 4:** Partitioned heritability for estimates for lung cancer

| Category | Pr(SNPs) | Pr(h2) | SE | Enrichment | SE | Enrichment P |
|---|---|---|---|---|---|---|
| CpG Content 50kb | 0.010 | 0.014 | 0.001 | 1.351 | 0.061 | 2.07E-07 |
| Background Selection Statistic | 0.178 | 0.251 | 0.019 | 1.412 | 0.108 | 1.03E-06 |
| SuperEnhancer Hnisz | 0.167 | 0.367 | 0.048 | 2.194 | 0.285 | 4.40E-06 |
| SuperEnhancer Hnisz (500bp window) | 0.170 | 0.350 | 0.051 | 2.052 | 0.298 | 7.61E-05 |
| Coding UCSC (500bp window) | 0.064 | 0.364 | 0.082 | 5.710 | 1.295 | 4.12E-04 |
| Intron UCSC (500bp window) | 0.397 | 0.574 | 0.055 | 1.447 | 0.137 | 1.11E-03 |
| H3K27ac PGC2 | 0.269 | 0.700 | 0.128 | 2.605 | 0.477 | 1.28E-03 |
| Conserved Vertebrate phastCons (500bp window) | 0.407 | 0.857 | 0.147 | 2.106 | 0.362 | 3.60E-03 |
| Conserved Mammal phastCons (500bp window) | 0.339 | 0.791 | 0.149 | 2.332 | 0.438 | 3.93E-03 |
| Conserved Primate phastCons (500bp window) | 0.176 | 0.524 | 0.130 | 2.981 | 0.742 | 5.34E-03 |
| Conserved LindbladToh (500bp window) | 0.330 | 0.761 | 0.144 | 2.302 | 0.436 | 5.36E-03 |
| Conserved LindbladToh | 0.026 | 0.322 | 0.110 | 12.552 | 4.268 | 0.011 |
| H3K27ac Hnisz (500bp window) | 0.420 | 0.670 | 0.095 | 1.594 | 0.227 | 0.012 |
| DHS Trynka (500bp window) | 0.496 | 0.901 | 0.157 | 1.817 | 0.316 | 0.013 |
| Conserved Primate phastCons | 0.019 | 0.228 | 0.086 | 11.826 | 4.454 | 0.013 |
| UTR 3 UCSC (500bp window) | 0.026 | 0.144 | 0.051 | 5.452 | 1.912 | 0.015 |
| TFBS ENCODE (500bp window) | 0.341 | 0.736 | 0.163 | 2.156 | 0.479 | 0.018 |
| Conserved Mammal phastCons | 0.021 | 0.447 | 0.172 | 20.855 | 8.021 | 0.022 |
| Intron UCSC | 0.387 | 0.506 | 0.062 | 1.307 | 0.160 | 0.041 |
| H3K9ac Trynka (500bp window) | 0.230 | 0.572 | 0.163 | 2.489 | 0.711 | 0.046 |
| Conserved Vertebrate phastCons | 0.029 | 0.400 | 0.174 | 13.592 | 5.905 | 0.046 |
| TSS Hoffman (500bp window) | 0.034 | 0.187 | 0.076 | 5.451 | 2.201 | 0.050 |
| H3K27ac Hnisz | 0.389 | 0.536 | 0.073 | 1.379 | 0.188 | 0.055 |
| DGF ENCODE (500bp window) | 0.538 | 0.813 | 0.140 | 1.510 | 0.259 | 0.058 |
| TFBS ENCODE | 0.131 | 0.489 | 0.188 | 3.728 | 1.430 | 0.064 |
| H3K4me1 Trynka | 0.424 | 0.705 | 0.158 | 1.664 | 0.372 | 0.084 |
| UTR 5 UCSC (500bp window) | 0.027 | 0.166 | 0.081 | 6.185 | 3.006 | 0.091 |
| H3K4me1 Trynka (500bp window) | 0.606 | 0.772 | 0.098 | 1.274 | 0.161 | 0.106 |
| Enhancer Hoffman (500bp window) | 0.090 | 0.225 | 0.086 | 2.502 | 0.955 | 0.114 |
| Fetal DHS Trynka (500bp window) | 0.283 | 0.557 | 0.172 | 1.967 | 0.606 | 0.121 |
| Transcr Hoffman | 0.346 | 0.571 | 0.159 | 1.651 | 0.460 | 0.125 |
| H3K27ac PGC2 (500bp window) | 0.335 | 0.507 | 0.119 | 1.512 | 0.356 | 0.126 |
| Promoter UCSC (500bp window) | 0.057 | 0.170 | 0.076 | 2.984 | 1.342 | 0.149 |
| BLUEPRINT DNA methylation MaxCPP | 0.032 | 0.137 | 0.072 | 4.306 | 2.271 | 0.150 |
| H3K4me3 Trynka (500bp window) | 0.255 | 0.441 | 0.135 | 1.729 | 0.530 | 0.175 |
| H3K4me3 Trynka | 0.133 | 0.293 | 0.120 | 2.204 | 0.901 | 0.181 |
| Synonymous | 0.003 | 0.068 | 0.049 | 21.735 | 15.804 | 0.190 |
| BLUEPRINT H3K27acQTL MaxCPP | 0.017 | 0.071 | 0.043 | 4.315 | 2.573 | 0.201 |

| | | | | | | |
|---|---|---|---|---|---|---|
| GTEx eQTL MaxCPP | 0.010 | 0.046 | 0.030 | 4.483 | 2.908 | 0.214 |
| Bivariate Flanking TSS/Enhancer | 0.014 | 0.096 | 0.070 | 7.050 | 5.164 | 0.243 |
| H3K9ac Trynka | 0.125 | 0.261 | 0.123 | 2.079 | 0.983 | 0.275 |
| GERP.RSsup4 | 0.008 | 0.068 | 0.056 | 8.287 | 6.909 | 0.299 |
| PromoterFlanking Hoffman | 0.008 | 0.070 | 0.060 | 8.483 | 7.258 | 0.306 |
| BLUEPRINT H3K4me1QTL MaxCPP | 0.013 | 0.047 | 0.034 | 3.482 | 2.515 | 0.320 |
| H3K4me3 peaks Trynka | 0.042 | 0.163 | 0.127 | 3.902 | 3.049 | 0.348 |
| Coding UCSC | 0.014 | 0.069 | 0.060 | 4.839 | 4.228 | 0.367 |
| Promoter UCSC | 0.046 | 0.154 | 0.124 | 3.329 | 2.680 | 0.387 |
| Enhancer Hoffman | 0.042 | 0.134 | 0.105 | 3.182 | 2.506 | 0.389 |
| WeakEnhancer Hoffman (500bp window) | 0.089 | 0.202 | 0.130 | 2.273 | 1.461 | 0.392 |
| Bivariate Flanking TSS/Enhancer (500bp window) | 0.031 | 0.084 | 0.065 | 2.682 | 2.091 | 0.416 |
| Enhancer Andersson (500bp window) | 0.019 | 0.055 | 0.057 | 2.882 | 2.991 | 0.534 |
| UTR 3 UCSC | 0.011 | 0.035 | 0.038 | 3.106 | 3.357 | 0.534 |
| TSS Hoffman | 0.018 | 0.058 | 0.071 | 3.267 | 4.006 | 0.573 |
| DGF ENCODE | 0.136 | 0.014 | 0.233 | 0.100 | 1.713 | 0.599 |
| PromoterFlanking Hoffman (500bp window) | 0.033 | 0.068 | 0.070 | 2.055 | 2.122 | 0.620 |
| FetalDHS Trynka | 0.084 | 0.004 | 0.172 | 0.042 | 2.049 | 0.633 |
| DHS Trynka | 0.166 | 0.074 | 0.223 | 0.445 | 1.342 | 0.669 |
| CTCF Hoffman (500bp window) | 0.071 | 0.143 | 0.179 | 2.017 | 2.522 | 0.681 |
| H3K4me1 peaks Trynka | 0.170 | 0.242 | 0.214 | 1.427 | 1.257 | 0.733 |
| H3K9ac peaks Trynka | 0.038 | 0.002 | 0.119 | 0.047 | 3.085 | 0.754 |
| Transcr Hoffman (500bp window) | 0.762 | 0.750 | 0.099 | 0.984 | 0.130 | 0.900 |

**Supplementary Table 5:** Additional 73 independent variants associated with pulmonary function identified and replicated in the UK Biobank cohort and used as genetic instruments in Mendelian randomization analyses of lung cancer risk

| CHR | Position (GRCh37) | SNP | Alleles | EAF | Discovery | | | Replication | Lead Phenotype[1] | INFO | Nearest Gene and Function | |
|---|---|---|---|---|---|---|---|---|---|---|---|---|
| | | | Effect/Other | | Beta | SE | P-value | P-value | | | | |
| 6 | 7146350 | rs4960289 | A/G | 0.427 | -0.0017 | 0.0002 | 9.6E-20 | 3.7E-14 | $FEV_1$/FVC | 0.986 | *RREB1* | intronic |
| 22 | 30598516 | rs6006399 | T/G | 0.884 | -0.0245 | 0.0028 | 1.9E-18 | 6.3E-05 | FEV1 | 1 | *HORMAD2* | intronic |
| 2 | 36724282 | rs1179500 | A/C | 0.721 | -0.0017 | 0.0002 | 3.6E-17 | 6.8E-05 | $FEV_1$/FVC | 0.988 | *CRIM1* | intronic |
| 16 | 67560613 | rs7196853 | T/C | 0.897 | -0.0025 | 0.0003 | 1.3E-16 | 1.4E-05 | $FEV_1$/FVC | 0.994 | *FAM65A* | intronic |
| 4 | 146087979 | rs116125427 | G/A | 0.922 | 0.0276 | 0.0034 | 2.3E-16 | 1.7E-06 | $FEV_1$ | 0.987 | *OTUD4* | intronic |
| 10 | 124230612 | rs12571363 | C/A | 0.893 | 0.0228 | 0.0028 | 3.2E-16 | 1.3E-06 | FVC | 0.995 | *HTRA1* | intronic |
| 8 | 103131300 | rs659398 | T/C | 0.269 | -0.0017 | 0.0002 | 4.9E-16 | 6.4E-05 | $FEV_1$/FVC | 0.983 | *NCALD* | intronic |
| 16 | 58062081 | rs41418849 | T/G | 0.940 | -0.0031 | 0.0004 | 5.2E-16 | 4.1E-06 | $FEV_1$/FVC | 0.949 | *MMP15* | intronic |
| 22 | 30254175 | rs9614084 | C/T | 0.521 | -0.0015 | 0.0002 | 7.2E-16 | 4.9E-11 | $FEV_1$/FVC | 0.996 | *ASCC2* | intergenic |
| 5 | 157022475 | rs72811372 | G/A | 0.895 | -0.0023 | 0.0003 | 2.3E-15 | 8.3E-06 | $FEV_1$/FVC | 0.998 | *AC008694.2* | intergenic |
| 1 | 221473248 | rs11118683 | C/T | 0.579 | -0.0140 | 0.0018 | 2.5E-15 | 2.7E-06 | FVC | 0.975 | *RP11-421L10.1* | intergenic |
| 14 | 54410919 | rs4444235 | T/C | 0.536 | 0.0014 | 0.0002 | 8.9E-15 | 4.4E-06 | $FEV_1$/FVC | 1 | *MIR5580* | intergenic |
| 5 | 120078424 | rs1125578 | A/C | 0.461 | -0.0014 | 0.0002 | 1.1E-14 | 1.3E-04 | $FEV_1$/FVC | 0.991 | *RNU4-69P* | intergenic |
| 19 | 46309203 | rs1548029 | A/C | 0.644 | -0.0015 | 0.0002 | 1.3E-14 | 5.5E-11 | $FEV_1$/FVC | 0.998 | *RSPH6A* | intronic |
| 2 | 42433247 | rs12466981 | C/T | 0.728 | -0.0015 | 0.0002 | 2.7E-14 | 5.9E-06 | $FEV_1$/FVC | 0.996 | *EML4* | intronic |
| 1 | 3425867 | rs2794359 | C/A | 0.895 | -0.0023 | 0.0003 | 2.8E-14 | 2.0E-10 | $FEV_1$/FVC | 0.986 | *MEGF6* | intronic |
| 5 | 158368797 | rs7443323 | T/A | 0.748 | 0.0151 | 0.0020 | 4.6E-14 | 1.6E-05 | FVC | 0.992 | *EBF1* | intronic |
| 5 | 43484257 | rs79904209 | C/T | 0.906 | 0.0229 | 0.0030 | 5.6E-14 | 2.2E-05 | $FEV_1$ | 1 | *TMEM267* | 5' UTR |
| 2 | 178210268 | rs6740092 | T/A | 0.145 | -0.0181 | 0.0025 | 1.9E-13 | 4.3E-05 | FVC | 0.992 | *LOC100130691* | ncRNA_intronic |
| 6 | 32652486 | rs9275156 | T/C | 0.766 | 0.0016 | 0.0002 | 2.4E-13 | 1.0E-05 | $FEV_1$/FVC | 0.998 | *HLA-DQB1* | intergenic |
| 18 | 20032854 | rs4800410 | A/C | 0.596 | -0.0134 | 0.0018 | 2.6E-13 | 1.3E-04 | $FEV_1$ | 0.988 | *RP11-863N1.4* | intergenic |
| 2 | 100568631 | rs11887136 | G/A | 0.868 | 0.0189 | 0.0026 | 7.6E-13 | 8.9E-05 | $FEV_1$ | 0.996 | *AFF3* | intronic |
| 6 | 152593102 | rs6904757 | A/G | 0.636 | 0.0129 | 0.0018 | 9.4E-13 | 1.1E-04 | FVC | 0.982 | *SYNE1* | intronic |
| 11 | 13166565 | rs11022690 | T/C | 0.545 | -0.0013 | 0.0002 | 1.2E-12 | 8.2E-06 | $FEV_1$/FVC | 0.995 | *RP11-413N13.1* | intergenic |
| 2 | 69407720 | rs10173269 | T/G | 0.539 | -0.0013 | 0.0002 | 1.3E-12 | 1.0E-04 | $FEV_1$/FVC | 1 | *ANTXR1* | intronic |

| | | | | | | | | | | | | |
|---|---|---|---|---|---|---|---|---|---|---|---|---|
| 6 | 33025953 | rs6457709 | A/G | 0.705 | 0.0014 | 0.0002 | 1.4E-12 | 5.3E-06 | $FEV_1$/FVC | 1 | *HLA-DPA1* | intergenic |
| 6 | 26207174 | rs9393688 | A/T | 0.738 | -0.0139 | 0.0020 | 1.7E-12 | 2.9E-05 | FVC | 0.992 | *HIST1H4E* | downstream |
| 6 | 19837774 | rs11759102 | C/T | 0.840 | -0.0171 | 0.0024 | 2.3E-12 | 5.2E-05 | $FEV_1$ | 0.989 | *ID4* | 5' UTR |
| 6 | 22072425 | rs196025 | A/G | 0.368 | -0.0130 | 0.0019 | 2.4E-12 | 9.0E-06 | $FEV_1$ | 0.985 | *CASC15* | ncRNA_intronic |
| 8 | 69570332 | rs17387279 | T/G | 0.819 | 0.0156 | 0.0023 | 5.0E-12 | 1.8E-04 | FVC | 0.990 | *C8orf34* | intronic |
| 1 | 92164100 | rs4233430 | C/T | 0.814 | -0.0016 | 0.0002 | 9.9E-12 | 1.4E-08 | $FEV_1$/FVC | 0.992 | *TGFBR3* | intronic |
| 15 | 77354847 | rs4886509 | C/A | 0.329 | -0.0013 | 0.0002 | 1.3E-11 | 1.0E-06 | $FEV_1$/FVC | 0.992 | *TSPAN3* | intronic |
| 17 | 60720058 | rs12452590 | T/G | 0.638 | 0.0013 | 0.0002 | 1.5E-11 | 5.8E-08 | $FEV_1$/FVC | 0.974 | *MRC2* | intronic |
| 5 | 170909410 | rs11745375 | C/T | 0.527 | -0.0012 | 0.0002 | 1.6E-11 | 1.3E-04 | $FEV_1$/FVC | 1 | *FGF18* | intergenic |
| 7 | 19050020 | rs28719767 | G/C | 0.711 | -0.0130 | 0.0020 | 5.1E-11 | 7.0E-05 | $FEV_1$ | 0.983 | *HDAC9* | intergenic |
| 22 | 18357509 | rs72490631 | T/C | 0.773 | 0.0140 | 0.0021 | 5.5E-11 | 2.0E-06 | $FEV_1$ | 0.994 | *MICAL3* | intronic |
| 2 | 230258493 | rs207672 | T/G | 0.340 | 0.0012 | 0.0002 | 7.2E-11 | 1.3E-07 | $FEV_1$/FVC | 0.992 | *DNER* | intronic |
| 11 | 12660894 | rs7927422 | T/C | 0.593 | 0.0115 | 0.0018 | 7.5E-11 | 2.1E-05 | FVC | 0.995 | *TEAD1* | intergenic |
| 3 | 73554922 | rs4677294 | T/A | 0.641 | 0.0121 | 0.0019 | 9.3E-11 | 1.6E-04 | $FEV_1$ | 0.989 | *PDZRN3* | intronic |
| 12 | 65962636 | rs10878300 | T/G | 0.198 | -0.0140 | 0.0022 | 1.3E-10 | 8.5E-05 | FVC | 0.992 | *LOC100507065* | ncRNA_intronic |
| 17 | 38489170 | rs2715554 | A/G | 0.849 | 0.0016 | 0.0003 | 1.5E-10 | 4.4E-05 | $FEV_1$/FVC | 1 | *RARA* | intronic |
| 16 | 88807608 | rs750739 | A/G | 0.833 | 0.0153 | 0.0024 | 1.8E-10 | 1.0E-04 | $FEV_1$ | 0.993 | *PIEZO1* | intronic |
| 1 | 92031492 | rs2125126 | G/A | 0.850 | -0.0016 | 0.0003 | 2.1E-10 | 8.3E-07 | $FEV_1$/FVC | 0.983 | *RP11-47K11.2* | intergenic |
| 5 | 132375622 | rs113638840 | A/G | 0.744 | 0.0130 | 0.0020 | 2.5E-10 | 2.0E-04 | $FEV_1$ | 0.996 | *HSPA4* | intergenic |
| 14 | 93507197 | rs1956028 | T/C | 0.874 | 0.0168 | 0.0027 | 4.3E-10 | 3.4E-07 | $FEV_1$ | 0.990 | *ITPK1* | intronic |
| 1 | 9498113 | rs9660890 | T/C | 0.803 | -0.0014 | 0.0002 | 5.8E-10 | 1.2E-04 | $FEV_1$/FVC | 0.992 | *RNA5SP40* | upstream |
| 6 | 90948093 | rs58453446 | G/C | 0.652 | -0.0015 | 0.0002 | 8.9E-10 | 3.5E-05 | $FEV_1$/FVC nvsmk | 0.998 | *BACH2* | intronic |
| 2 | 169479763 | rs10184235 | A/G | 0.753 | -0.0013 | 0.0002 | 1.1E-09 | 1.1E-06 | $FEV_1$/FVC | 0.999 | *CERS6* | intronic |
| 5 | 173303392 | rs55993676 | G/T | 0.713 | -0.0012 | 0.0002 | 1.4E-09 | 7.4E-05 | $FEV_1$/FVC | 0.998 | *CPEB4* | intergenic |
| 4 | 174582067 | rs10005540 | C/T | 0.386 | -0.0107 | 0.0018 | 2.6E-09 | 1.7E-04 | FVC | 0.978 | *RANP6* | intergenic |
| 1 | 22653424 | rs4233284 | C/G | 0.672 | 0.0109 | 0.0018 | 3.2E-09 | 1.3E-04 | FVC | 0.993 | *RP11-415K20.1* | intergenic |
| 12 | 28840892 | rs12313454 | A/G | 0.882 | -0.0158 | 0.0027 | 3.5E-09 | 9.2E-05 | FVC | 0.996 | *CCDC91* | intergenic |
| 8 | 64806567 | rs1425794 | G/T | 0.565 | -0.0103 | 0.0017 | 4.2E-09 | 1.5E-04 | FVC | 0.995 | *RP11-32K4.1* | ncRNA_intronic |

| | | | | | | | | | | | | |
|---|---|---|---|---|---|---|---|---|---|---|---|---|
| 7 | 140560023 | rs13227429 | T/C | 0.436 | 0.0102 | 0.0017 | 5.6E-09 | 6.6E-05 | FVC | 0.994 | *BRAF* | intronic |
| 2 | 67052289 | rs17032590 | A/G | 0.791 | -0.0013 | 0.0002 | 8.1E-09 | 8.0E-06 | $FEV_1$/FVC | 0.996 | *AC009474.1* | intergenic |
| 8 | 13194983 | rs1528624 | T/G | 0.505 | 0.0010 | 0.0002 | 8.4E-09 | 7.6E-09 | $FEV_1$/FVC | 0.990 | *DLC1* | intronic |
| 10 | 63839417 | rs4948502 | T/C | 0.573 | 0.0101 | 0.0018 | 8.9E-09 | 3.0E-07 | FVC | 0.995 | *ARID5B* | intronic |
| 5 | 148203236 | rs35684381 | T/C | 0.854 | -0.0015 | 0.0003 | 8.9E-09 | 3.4E-06 | $FEV_1$/FVC | 0.979 | *ADRB2* | intergenic |
| 4 | 145557467 | rs35797611 | T/C | 0.968 | -0.0029 | 0.0005 | 9.6E-09 | 1.9E-05 | $FEV_1$/FVC | 1 | *HHIP-AS1* | intergenic |
| 20 | 3660789 | rs603112 | T/C | 0.157 | -0.0014 | 0.0002 | 1.2E-08 | 1.3E-04 | $FEV_1$/FVC | 1 | *ADAM33* | intronic |
| 3 | 11642114 | rs1561073 | T/A | 0.262 | -0.0012 | 0.0002 | 1.6E-08 | 2.2E-05 | $FEV_1$/FVC | 0.992 | *VGLL4* | intronic |
| 15 | 67464291 | rs72743477 | A/G | 0.783 | 0.0012 | 0.0002 | 1.7E-08 | 1.1E-04 | $FEV_1$/FVC | 0.985 | *SMAD3* | intronic |
| 7 | 134567104 | rs28517513 | G/T | 0.751 | 0.0012 | 0.0002 | 2.1E-08 | 5.7E-05 | $FEV_1$/FVC | 0.990 | *CALD1* | intronic |
| 11 | 65324276 | rs11227223 | C/T | 0.945 | -0.0022 | 0.0004 | 2.2E-08 | 3.4E-07 | $FEV_1$/FVC | 0.995 | *LTBP3* | intronic |
| 17 | 13416372 | rs1978218 | G/T | 0.404 | -0.0132 | 0.0024 | 2.3E-08 | 1.4E-04 | FVC-nvsmk | 0.986 | *HS3ST3A1* | intronic |
| 4 | 90036240 | rs17821105 | A/T | 0.830 | 0.0013 | 0.0002 | 2.5E-08 | 2.3E-06 | $FEV_1$/FVC | 0.997 | *TIGD2* | downstream |
| 16 | 31030344 | rs4889526 | C/A | 0.625 | 0.0102 | 0.0018 | 2.9E-08 | 9.5E-09 | $FEV_1$ | 0.998 | *STX1B* | intergenic |
| 3 | 53672471 | rs9819463 | T/C | 0.795 | 0.0123 | 0.0022 | 2.9E-08 | 2.5E-07 | $FEV_1$ | 0.994 | *CACNA1D* | intronic |
| 9 | 129416317 | rs10987386 | C/T | 0.813 | 0.0013 | 0.0002 | 3.3E-08 | 1.2E-08 | $FEV_1$/FVC | 0.982 | *LMX1B* | intronic |
| 2 | 43762112 | rs77972916 | G/A | 0.924 | -0.0019 | 0.0003 | 3.3E-08 | 3.0E-05 | $FEV_1$/FVC | 0.995 | *THADA* | intronic |
| 18 | 38023468 | rs4636990 | G/A | 0.457 | -0.0096 | 0.0017 | 3.8E-08 | 7.3E-05 | FVC | 0.990 | *RNU7-145P* | intergenic |
| 8 | 109446828 | rs10089406 | T/C | 0.404 | 0.0010 | 0.0002 | 4.4E-08 | 5.0E-06 | $FEV_1$/FVC | 0.987 | *EIF3E* | intergenic |

1. Lead phenotype refers to the lung function trait for which this variant was selected to be a genetic instrument. For each SNP the lead phenotype has the smallest p-value.

**Supplementary Table 6:** Descriptive characteristics of the OncoArray lung cancer population

| | | Lung Cancer Cases | | Controls | |
|---|---|---|---|---|---|
| | | Number | (%) | Number | (%) |
| Age | ≤50 years | 3112 | (12) | 6,032 | (12) |
| | >50 years | 23025 | (88) | 44,075 | (88) |
| Sex | Male | 18208 | (62) | 27,178 | (53) |
| | Female | 11059 | (38) | 24,069 | (47) |
| Smoking status | Never smokers | 2355 | (9) | 7,504 | (31) |
| | Ever smokers | 23223 | (91) | 16,964 | (69) |
| | Former smokers | 9037 | (35) | 8,554 | (35) |
| | Current smokers | 13356 | (52) | 7,477 | (31) |
| Histology | Adenocarcinoma | 11273 | (39) | | |
| | Squamous cell carcinoma | 7426 | (35) | | |
| | Small cell carcinoma | 2664 | (9) | | |
| Total | | 29266 | (100) | 56,450 | (100) |

**Supplementary Table 7:** Mendelian randomization (MR) odds ratio (OR) estimates for lung cancer overall and by histology for a genetically predicted 1-SD decrease in a standardized $FEV_1$ z-score

| Outcome | MR Estimator | $N_{SNPs}$ | OR | (95% CI) | P |
|---|---|---|---|---|---|
| Lung Cancer (cases: n=29,266 controls: n=56,450) | IVW-RE | 193 | 1.27 | (1.01 – 1.60) | 0.041 |
| | Maximum likelihood | 193 | 1.28 | (1.12 – 1.47) | $3.4\times10^{-4}$ |
| | Weighted median | 193 | 1.06 | (0.86 – 1.32) | 0.572 |
| | MR RAPS | 193 | 1.13 | (0.91 – 1.40) | 0.255 |
| | MR Egger slope | 193 | 1.65 | (0.70 – 3.87) | 0.251 |
| | Outliers filtered: | | | | |
| | IVW-RE | 157 | 1.12 | (0.97 – 1.30) | 0.132 |
| | Maximum likelihood | 157 | 1.12 | (0.97 – 1.30) | 0.132 |
| | Weighted median | 157 | 1.07 | (0.86 – 1.32) | 0.554 |
| | MR RAPS | 157 | 1.09 | (0.93 – 1.28) | 0.299 |
| | MR Egger slope | 157 | 0.84 | (0.47 – 1.50) | 0.555 |
| Adenocarcinoma (cases: n=11,273 controls: n=56,450) | IVW-RE | 192 | 0.99 | (0.79 – 1.26) | 0.965 |
| | Maximum likelihood | 192 | 0.99 | (0.83 – 1.19) | 0.956 |
| | Weighted median | 192 | 1.07 | (0.81 – 1.43) | 0.633 |
| | MR RAPS | 192 | 1.00 | (0.79 – 1.27) | 0.987 |
| | MR Egger slope | 192 | 0.91 | (0.39 – 2.10) | 0.827 |
| | Outliers filtered: | | | | |
| | IVW-RE | 169 | 1.03 | (0.86 – 1.24) | 0.714 |
| | Maximum likelihood | 169 | 1.03 | (0.85 – 1.26) | 0.728 |
| | Weighted median | 169 | 1.08 | (0.81 – 1.44) | 0.612 |
| | MR RAPS | 169 | 1.05 | (0.86 – 1.29) | 0.634 |
| | MR Egger slope | 169 | 0.80 | (0.39 – 1.66) | 0.557 |
| Squamous carcinoma (cases: n=7426) controls: n=56,450) | IVW-RE | 190 | 2.03 | (1.43 – 2.86) | $6.3\times10^{-5}$ |
| | Maximum likelihood | 190 | 2.04 | (1.64 – 2.54) | $1.2\times10^{-10}$ |
| | Weighted median | 190 | 1.54 | (1.09 – 2.17) | 0.014 |
| | MR RAPS | 190 | 1.81 | (1.30 – 2.50) | $3.7\times10^{-4}$ |
| | MR Egger slope | 190 | 5.43 | (1.50 – 19.63) | 0.011 |
| | Outliers filtered: | | | | |
| | IVW-RE | 156 | 1.51 | (1.21 – 1.88) | $2.2\times10^{-4}$ |
| | Maximum likelihood | 156 | 1.50 | (1.19 – 1.90) | $6.7\times10^{-4}$ |
| | Weighted median | 156 | 1.44 | (1.02 – 2.04) | 0.040 |
| | MR RAPS | 156 | 1.48 | (1.16 – 1.88) | $1.7\times10^{-3}$ |
| | MR Egger slope | 156 | 2.11 | (0.89 – 5.02) | 0.092 |

IVW – RE : Inverse variance weighted – multiplicative random effects
RAPS : Robust adjusted profile score – Huber loss function
PRESSO : Randomization Pleiotropy RESidual Sum and Outlier

**Supplementary Table 8:** Mendelian randomization (MR) odds ratio (OR) estimates for the risk of lung cancer overall and by histology for a genetically predicted 1-SD decrease in a standardized FVC z-score.

| Outcome | MR Estimator | $N_{SNPs}$ | OR | (95% CI) | P |
|---|---|---|---|---|---|
| Lung Cancer (cases: n=29,266 controls: n=56,450) | IVW-RE | 144 | 1.16 | (0.91 – 1.47) | 0.232 |
| | Maximum likelihood | 144 | 1.16 | (0.98 – 1.37) | 0.078 |
| | Weighted median | 144 | 1.05 | (0.81 – 1.35) | 0.726 |
| | MR RAPS | 144 | 1.09 | (0.87 – 1.36) | 0.445 |
| | MR Egger slope | 144 | 0.75 | (0.28 – 2.04) | 0.576 |
| | Outliers filtered: | | | | |
| | IVW-RE | 124 | 1.14 | (0.97 – 1.34) | 0.121 |
| | Maximum likelihood | 124 | 1.14 | (0.96 – 1.37) | 0.144 |
| | Weighted median | 124 | 1.05 | (0.81 – 1.37) | 0.699 |
| | MR RAPS | 124 | 1.13 | (0.94 – 1.36) | 0.206 |
| | MR Egger slope | 124 | 0.45 | (0.20 – 1.05) | 0.067 |
| Adenocarcinoma (cases: n=11,273 controls: n=56,450) | IVW-RE | 143 | 0.92 | (0.68 – 1.24) | 0.576 |
| | Maximum likelihood | 143 | 0.92 | (0.73 – 1.16) | 0.462 |
| | Weighted median | 143 | 1.03 | (0.72 – 1.46) | 0.878 |
| | MR RAPS | 143 | 0.94 | (0.70 – 1.26) | 0.689 |
| | MR Egger slope | 143 | 0.64 | (0.19 – 2.18) | 0.478 |
| | Outliers filtered: | | | | |
| | IVW-RE | 124 | 0.96 | (0.76 – 1.22) | 0.744 |
| | Maximum likelihood | 124 | 0.96 | (0.75 – 1.23) | 0.760 |
| | Weighted median | 124 | 1.04 | (0.72 – 1.49) | 0.842 |
| | MR RAPS | 124 | 0.98 | (0.76 – 1.27) | 0.904 |
| | MR Egger slope | 124 | 0.92 | (0.29 – 2.91) | 0.884 |
| Squamous carcinoma (cases: n=7426) controls: n=56,450) | IVW-RE | 140 | 1.67 | (1.13 – 2.46) | 0.011 |
| | Maximum likelihood | 140 | 1.68 | (1.28 – 2.20) | $1.8\times10^{-4}$ |
| | Weighted median | 140 | 1.19 | (0.81 – 1.76) | 0.381 |
| | MR RAPS | 140 | 1.46 | (1.11 – 1.92) | $6.5\times10^{-3}$ |
| | MR Egger slope | 140 | 0.73 | (0.14 – 3.70) | 0.700 |
| | Outliers filtered: | | | | |
| | IVW | 122 | 1.27 | (0.96 – 1.66) | 0.092 |
| | Maximum likelihood | 122 | 1.27 | (0.95 – 1.69) | 0.103 |
| | Weighted median | 122 | 1.18 | (0.78 – 1.78) | 0.429 |
| | MR RAPS | 122 | 1.25 | (0.93 – 1.69) | 0.139 |
| | MR Egger slope | 122 | 0.43 | (0.13 – 1.41) | 0.167 |

IVW – RE : Inverse variance weighted – multiplicative random effects
RAPS : Robust adjusted profile score – Huber loss function

**Supplementary Table 9:** Mendelian randomization (MR) odds ratio (OR) estimates for the risk of lung cancer overall and by histology for a genetically predicted 10% decrease in $FEV_1$/FVC.

| Outcome | MR Estimator | $N_{SNPs}$ | OR | (95% CI) | P |
|---|---|---|---|---|---|
| Lung Cancer (cases: n=29,266 controls: n=56,450) | IVW-RE | 264 | 1.18 | (1.01 – 1.38) | 0.035 |
| | Maximum likelihood | 264 | 1.18 | (1.07 – 1.31) | $1.6\times10^{-3}$ |
| | Weighted median | 264 | 1.10 | (0.92 – 1.30) | 0.296 |
| | MR RAPS | 264 | 1.11 | (0.97 – 1.29) | 0.137 |
| | MR Egger slope | 264 | 1.15 | (0.77 – 1.71) | 0.485 |
| | Outliers filtered: | | | | |
| | IVW-RE | 226 | 1.10 | (0.99 – 1.22) | 0.082 |
| | Maximum likelihood | 226 | 1.10 | (0.98 – 1.23) | 0.100 |
| | Weighted median | 226 | 1.12 | (0.93 – 1.34) | 0.237 |
| | MR RAPS | 226 | 1.09 | (0.97 – 1.23) | 0.143 |
| | MR Egger slope | 255 | 1.22 | (0.91 – 1.64) | 0.184 |
| Adenocarcinoma (cases: n=11,273 controls: n=56,450) | IVW-RE | 265 | 1.11 | (0.93 – 1.32) | 0.245 |
| | Maximum likelihood | 265 | 1.11 | (0.97 – 1.27) | 0.145 |
| | Weighted median | 265 | 1.16 | (0.91 – 1.46) | 0.226 |
| | MR RAPS | 265 | 1.12 | (0.94 – 1.33) | 0.207 |
| | MR Egger slope | 265 | 1.29 | (0.84 – 1.98) | 0.240 |
| | Outliers filtered: | | | | |
| | IVW-RE | 239 | 1.16 | (1.01 – 1.34) | 0.032 |
| | Maximum likelihood | 239 | 1.17 | (1.01 – 1.35) | 0.039 |
| | Weighted median | 239 | 1.23 | (1.00 – 1.53) | 0.054 |
| | MR RAPS | 239 | 1.18 | (1.02 – 1.38) | 0.030 |
| | MR Egger slope | 239 | 1.43 | (1.00 – 2.05) | 0.052 |
| Squamous carcinoma (cases: n=7426) controls: n=56,450) | IVW-RE | 263 | 1.30 | (1.04 – 1.62) | 0.022 |
| | Maximum likelihood | 263 | 1.29 | (1.10 – 1.53) | $2.4\times10^{-3}$ |
| | Weighted median | 263 | 1.04 | (0.79 – 1.36) | 0.775 |
| | MR RAPS | 263 | 1.13 | (0.91 – 1.40) | 0.255 |
| | MR Egger slope | 263 | 1.25 | (0.70 – 2.24) | 0.446 |
| | Outliers filtered: | | | | |
| | IVW-RE | 239 | 1.13 | (0.95 – 1.34) | 0.164 |
| | Maximum likelihood | 239 | 1.13 | (0.95 – 1.34) | 0.173 |
| | Weighted median | 239 | 1.04 | (0.79 – 1.37) | 0.785 |
| | MR RAPS | 239 | 1.09 | (0.90 – 1.31) | 0.372 |
| | MR Egger slope | 239 | 1.03 | (0.65 – 1.62) | 0.912 |

IVW – RE : Inverse variance weighted – multiplicative random effects
RAPS : Robust adjusted profile score – Huber loss function

**Supplementary Table 10:** Mendelian randomization (MR) odds ratio (OR) estimates for risk of lung cancer in never smokers (2355 cases, 7504 controls) corresponding to a 1-SD decrease in $FEV_1$ and FVC z-scores and 10% decrease in $FEV_1$/FVC. Genetic instruments for pulmonary function were developed in a separate GWAS of never smokers in the UK Biobank.

| Phenotype | MR Estimator | $N_{SNPs}$ | OR | (95% CI) | P |
|---|---|---|---|---|---|
| | IVW-RE | 76 | 0.99 | (0.55 – 1.80) | 0.984 |
| | Maximum likelihood | 76 | 0.99 | (0.59 – 1.68) | 0.982 |
| | Weighted median | 76 | 0.75 | (0.36 – 1.58) | 0.450 |
| | MR RAPS | 76 | 1.02 | (0.55 – 1.88) | 0.958 |
| | MR Egger slope | 76 | 0.87 | (0.08 – 9.65) | 0.912 |
| $FEV_1$ | Outliers filtered: | | | | |
| | IVW-RE | 68 | 1.11 | (0.69 – 1.80) | 0.669 |
| | Maximum likelihood | 68 | 1.11 | (0.64 – 1.93) | 0.704 |
| | Weighted median | 68 | 0.81 | (0.37 – 1.79) | 0.606 |
| | MR RAPS | 68 | 1.13 | (0.64 – 2.00) | 0.674 |
| | MR Egger slope | 68 | 1.29 | (0.15 – 11.07) | 0.816 |
| | IVW-RE | 57 | 0.79 | (0.37 – 1.66) | 0.527 |
| | Maximum likelihood | 57 | 0.78 | (0.41 – 1.49) | 0.452 |
| | Weighted median | 57 | 0.54 | (0.21 – 1.41) | 0.211 |
| | MR RAPS | 57 | 0.66 | (0.30 – 1.44) | 0.296 |
| | MR Egger slope | 57 | 0.58 | (0.02 – 18.48) | 0.758 |
| FVC | Outliers filtered: | | | | |
| | IVW-RE | 52 | 0.60 | (0.33 – 1.10) | 0.101 |
| | Maximum likelihood | 52 | 0.59 | (0.30 – 1.17) | 0.130 |
| | Weighted median | 52 | 0.52 | (0.20 – 1.36) | 0.181 |
| | MR RAPS | 52 | 0.57 | (0.28 – 1.15) | 0.115 |
| | MR Egger slope | 52 | 0.42 | (0.02 – 7.83) | 0.565 |
| | IVW-RE | 112 | 1.60 | (1.05 – 2.45) | 0.030 |
| | Maximum likelihood | 112 | 1.61 | (1.10 – 2.35) | 0.014 |
| | Weighted median | 112 | 1.41 | (0.78 – 2.57) | 0.254 |
| | MR RAPS | 112 | 1.57 | (1.03 – 2.37) | 0.035 |
| | MR Egger slope | 112 | 0.86 | (0.25 – 2.99) | 0.809 |
| $FEV_1$/FVC | Outliers filtered: | | | | |
| | IVW-RE | 103 | 1.55 | (1.09 – 2.19) | 0.014 |
| | Maximum likelihood | 103 | 1.56 | (1.05 – 2.30) | 0.027 |
| | Weighted median | 103 | 1.47 | (0.83 – 2.61) | 0.186 |
| | MR RAPS | 103 | 1.54 | (1.03 – 2.32) | 0.035 |
| | MR Egger slope | 103 | 1.50 | (0.47 – 4.83) | 0.494 |

IVW – RE : Inverse variance weighted – multiplicative random effects
RAPS : Robust adjusted profile score – Huber loss function

**Supplementary Table 11:** Mendelian randomization (MR) odds ratio (OR) estimates for risk of lung cancer in smokers (23223 cases, 16964 controls) corresponding to a 1-SD decrease in $FEV_1$ and FVC z-scores and a 10% decrease in $FEV_1$/FVC. Genetic instruments for pulmonary function were not specific to smokers.

| Phenotype | MR Estimator | $N_{SNPs}$ | OR | (95% CI) | P |
|---|---|---|---|---|---|
| | IVW-RE | 188 | 1.26 | (0.98 – 1.63) | 0.075 |
| | Maximum likelihood | 188 | 1.26 | (1.06 – 1.50) | $9.1\times10^{-3}$ |
| | Weighted median | 188 | 1.12 | (0.85 – 1.47) | 0.433 |
| | MR RAPS | 188 | 1.16 | (0.90 – 1.48) | 0.246 |
| | MR Egger slope | 188 | 1.76 | (0.69 – 4.51) | 0.239 |
| $FEV_1$ | Outliers filtered: | | | | |
| | IVW-RE | 163 | 1.09 | (0.91 – 1.31) | 0.363 |
| | Maximum likelihood | 163 | 1.09 | (0.90 – 1.31) | 0.368 |
| | Weighted median | 163 | 1.06 | (0.81 – 1.39) | 0.666 |
| | MR RAPS | 163 | 1.05 | (0.86 – 1.29) | 0.600 |
| | MR Egger slope | 163 | 0.81 | (0.41 – 1.61) | 0.551 |
| | IVW-RE | 139 | 1.20 | (0.91 – 1.58) | 0.206 |
| | Maximum likelihood | 139 | 1.20 | (0.97 – 1.48) | 0.100 |
| | Weighted median | 139 | 1.10 | (0.80 – 1.53) | 0.546 |
| | MR RAPS | 139 | 1.10 | (0.85 – 1.43) | 0.472 |
| | MR Egger slope | 139 | 1.05 | (0.33 – 3.32) | 0.935 |
| FVC | Outliers filtered: | | | | |
| | IVW-RE | 124 | 1.09 | (0.88 – 1.33) | 0.434 |
| | Maximum likelihood | 124 | 1.09 | (0.87 – 1.36) | 0.465 |
| | Weighted median | 124 | 1.07 | (0.77 – 1.48) | 0.685 |
| | MR RAPS | 124 | 1.08 | (0.86 – 1.36) | 0.525 |
| | MR Egger slope | 124 | 0.62 | (0.24 – 1.61) | 0.329 |
| | IVW-RE | 258 | 1.21 | (1.01 – 1.46) | 0.039 |
| | Maximum likelihood | 258 | 1.21 | (1.06 – 1.39) | $4.5\times10^{-3}$ |
| | Weighted median | 258 | 0.96 | (0.78 – 1.19) | 0.711 |
| | MR RAPS | 258 | 1.13 | (0.94 – 1.35) | 0.185 |
| | MR Egger slope | 258 | 0.92 | (0.57 – 1.48) | 0.722 |
| $FEV_1$/FVC | Outliers filtered: | | | | |
| | IVW-RE | 223 | 1.15 | (1.01 – 1.31) | 0.038 |
| | Maximum likelihood | 223 | 1.15 | (0.99 – 1.33) | 0.062 |
| | Weighted median | 223 | 1.08 | (0.87 – 1.34) | 0.488 |
| | MR RAPS | 223 | 1.14 | (0.98 – 1.32) | 0.094 |
| | MR Egger slope | 223 | 1.01 | (0.67 – 1.51) | 0.972 |

IVW – RE : Inverse variance weighted – multiplicative random effects
RAPS : Robust adjusted profile score – Huber loss function

**Supplementary Table 12:** Summary of diagnostic tests conducted for Mendelian Randomization analyses

| Phenotype Outcome Pair | MR Egger | | Modified Cochran's Q | | $I^2_{GX}$[1] | Mean F | MR Steiger Test[2] | | Number of Outliers |
|---|---|---|---|---|---|---|---|---|---|
| | $\beta_{0\ Egger}$ | p-value | Q statistic | p-value | | | Ratio | p-value | |
| $FEV_1$ Lung cancer | -0.004 | 0.53 | 585.58 | $2.1\times10^{-41}$ | 0.981 | 52.09 | 18.24 | $<2\times10^{-16}$ | 36 |
| $FEV_1$ Adenocarcinoma | 0.001 | 0.83 | 326.84 | $3.4\times10^{-9}$ | 0.981 | 53.54 | 17.01 | $<2\times10^{-16}$ | 23 |
| $FEV_1$ Squamous | -0.015 | 0.12 | 505.33 | $1.1\times10^{-30}$ | 0.981 | 52.13 | 11.20 | $<2\times10^{-16}$ | 34 |
| $FEV_1$/FVC Lung cancer | $4.4\times10^{-4}$ | 0.90 | 602.10 | $1.2\times10^{-28}$ | 0.984 | 63.25 | 43.25 | $<2\times10^{-16}$ | 38 |
| $FEV_1$/FVC Adenocarcinoma | -0.003 | 0.44 | 419.6 | $3.4\times10^{-9}$ | 0.985 | 66.11 | 38.65 | $<2\times10^{-16}$ | 26 |
| $FEV_1$/FVC Squamous | 0.001 | 0.90 | 479.9 | $5.3\times10^{-15}$ | 0.984 | 62.60 | 28.33 | $<2\times10^{-16}$ | 24 |
| FVC Lung cancer | 0.006 | 0.38 | 310.4 | $2.1\times10^{-14}$ | 0.979 | 48.02 | 13.60 | $<2\times10^{-16}$ | 20 |
| FVC Adenocarcinoma | 0.005 | 0.55 | 240.2 | $5.0\times10^{-7}$ | 0.979 | 47.48 | 9.99 | $<2\times10^{-16}$ | 19 |
| FVC Squamous | 0.012 | 0.30 | 304.4 | $2.2\times10^{-14}$ | 0.979 | 48.29 | 7.62 | $<2\times10^{-16}$ | 18 |
| $FEV_1$ Never smokers | 0.002 | 0.91 | 98.54 | 0.036 | 0.977 | 45.03 | 3.87 | $3.2\times10^{-7}$ | 8 |
| $FEV_1$/FVC Never smokers | 0.014 | 0.30 | 142.82 | 0.023 | 0.983 | 59.38 | 10.58 | $<2\times10^{-16}$ | 9 |
| FVC Never smokers | 0.005 | 0.86 | 78.07 | 0.027 | 0.976 | 40.82 | 2.22 | $4.1\times10^{-4}$ | 5 |
| $FEV_1$ Smokers | -0.005 | 0.47 | 419.05 | $7.7\times10^{-17}$ | 0.981 | 52.02 | 12.19 | $<2\times10^{-16}$ | 25 |
| $FEV_1$/FVC Smokers | 0.005 | 0.22 | 490.72 | $6.0\times10^{-17}$ | 0.984 | 62.13 | 27.12 | $<2\times10^{-16}$ | 35 |
| FVC Smokers | 0.002 | 0.82 | 239.11 | $2.1\times10^{-7}$ | 0.979 | 48.77 | 8.76 | $<2\times10^{-16}$ | |
| After outlier filtering: | | | | | | | | | |
| $FEV_1$ Lung cancer | 0.004 | 0.36 | 153.53 | 0.52 | 0.980 | 50.46 | 21.47 | $<2\times10^{-16}$ | |
| $FEV_1$ Adenocarcinoma | 0.004 | 0.48 | 153.06 | 0.79 | 0.981 | 52.65 | 18.77 | $<2\times10^{-16}$ | |
| $FEV_1$ Squamous | -0.004 | 0.50 | 141.12 | 0.80 | 0.981 | 52.55 | 14.09 | $<2\times10^{-16}$ | |
| $FEV_1$/FVC Lung cancer | -0.002 | 0.44 | 200.22 | 0.88 | 0.984 | 61.29 | 47.97 | $<2\times10^{-16}$ | |
| $FEV_1$/FVC Adenocarcinoma | -0.004 | 0.22 | 219.37 | 0.80 | 0.985 | 65.64 | 40.71 | $<2\times10^{-16}$ | |
| $FEV_1$/FVC Squamous | 0.002 | 0.66 | 228.8 | 0.64 | 0.984 | 61.47 | 30.67 | $<2\times10^{-16}$ | |
| FVC Lung cancer | 0.013 | 0.03 | 106.2 | 0.86 | 0.979 | 46.83 | 16.09 | $<2\times10^{-16}$ | |
| FVC Adenocarcinoma | 0.001 | 0.93 | 111.2 | 0.77 | 0.978 | 46.04 | 10.81 | $<2\times10^{-16}$ | |
| FVC Squamous | 0.014 | 0.10 | 110.4 | 0.72 | 0.979 | 47.99 | 9.23 | $<2\times10^{-16}$ | |
| $FEV_1$ Never smokers | -0.003 | 0.89 | 52.68 | 0.90 | 0.978 | 45.40 | 4.49 | $2.4\times10^{-12}$ | |
| $FEV_1$/FVC Never smokers | 0.001 | 0.96 | 81.84 | 0.93 | 0.983 | 59.64 | 11.56 | $<2\times10^{-16}$ | |
| FVC Never smokers | 0.001 | 0.97 | 45.45 | 0.73 | 0.976 | 40.40 | 2.45 | $2.7\times10^{-7}$ | |
| $FEV_1$ Smokers | -0.005 | 0.39 | 160.75 | 0.52 | 0.980 | 50.70 | 13.92 | $<2\times10^{-16}$ | |
| $FEV_1$/FVC Smokers | 0.002 | 0.49 | 181.48 | 0.98 | 0.983 | 59.27 | 29.55 | $<2\times10^{-16}$ | |
| FVC Smokers | 0.008 | 0.24 | 105.00 | 0.88 | 0.979 | 48.74 | 10.25 | $<2\times10^{-16}$ | |

1. Values close to 1 indicate that the no measurement error (NOME) assumption is unlikely to be violated. Low $I^2_{GX}$ estimates, less than 0.9, suggest that inferences should be interpreted with caution and adjustment methods should be considered
2. A higher ratio value indicates that the inferred direction is less susceptible to measurement error

**Supplementary Table 13:** Odds ratio (OR) estimates for a genetically predicted 1-SD decrease in standardized $FEV_1$ z-scores and 10% decrease in $FEV_1$/FVC after manual filtering to remove instruments associated with smoking status (ever/never), cigarette pack-years, or bodyfat percentage ($p<1\times10^{-5}$).

| MR Estimator | | | | | | | | |
|---|---|---|---|---|---|---|---|---|
| | $FEV_1$ and Lung Cancer | | | | | | | |
| | Manually filtered | | | | Manual filter and Outlier detection | | | |
| | $N_{SNPs}$ | OR | (95% CI) | P-value | $N_{SNPs}$ | OR | (95% CI) | P-value |
| IVW-RE | 168 | 1.25 | (0.98 - 1.59) | 0.067 | 141 | 1.08 | (0.93 - 1.26) | 0.320 |
| Maximum likelihood | 168 | 1.25 | (1.08 - 1.45) | 0.002 | 141 | 1.08 | (0.93 - 1.27) | 0.321 |
| Weighted median | 168 | 1.05 | (0.84 - 1.31) | 0.661 | 141 | 1.04 | (0.82 - 1.32) | 0.766 |
| MR RAPS | 168 | 1.11 | (0.89 - 1.38) | 0.364 | 141 | 1.05 | (0.89 - 1.25) | 0.564 |
| Egger intercept | 168 | -0.011 | | 0.091 | 141 | 0.002 | | 0.699 |
| Q statistic | 168 | 484.7 | | $1.0\times10^{-32}$ | 141 | 141.0 | | 0.46 |
| | $FEV_1$ and Squamous Carcinoma | | | | | | | |
| | Manually filtered | | | | Manual filter and Outlier detection | | | |
| | $N_{SNPs}$ | OR | (95% CI) | P-value | $N_{SNPs}$ | OR | (95% CI) | P-value |
| IVW-RE | 165 | 2.02 | (1.40 - 2.92) | $1.9\times10^{-4}$ | 137 | 1.46 | (1.16 - 1.84) | 0.001 |
| Maximum likelihood | 165 | 2.02 | (1.60 - 2.55) | $3.1\times10^{-9}$ | 137 | 1.46 | (1.14 - 1.87) | 0.003 |
| Weighted median | 165 | 1.47 | (1.03 - 2.11) | 0.035 | 137 | 1.36 | (0.93 - 2.00) | 0.111 |
| MR RAPS | 165 | 1.81 | (1.29 - 2.56) | $7.1\times10^{-4}$ | 137 | 1.41 | (1.09 - 1.83) | 0.009 |
| Egger intercept | 165 | -0.02 | | 0.048 | 137 | -0.007 | | 0.280 |
| MR Egger slope | 165 | 7.56 | (1.97 – 29.04) | 0.004 | 137 | 2.38 | (0.95 – 5.94) | 0.066 |
| Q statistic | 165 | 435.3 | | $1.9\times10^{-26}$ | 137 | 120.4 | | 0.827 |
| | $FEV_1$/FVC and Lung Cancer | | | | | | | |
| | Manually filtered | | | | Manual filter and Outlier detection | | | |
| | $N_{SNPs}$ | OR | (95% CI) | P-value | $N_{SNPs}$ | OR | (95% CI) | P-value |
| IVW-RE | 246 | 1.19 | (1.02 - 1.38) | 0.03 | 214 | 1.09 | (0.98 - 1.22) | 0.103 |
| Maximum likelihood | 246 | 1.19 | (1.07 - 1.32) | 0.001 | 214 | 1.09 | (0.97 - 1.22) | 0.133 |
| Weighted median | 246 | 1.11 | (0.93 - 1.33) | 0.25 | 214 | 1.15 | (0.96 - 1.39) | 0.130 |
| MR RAPS | 246 | 1.12 | (0.97 - 1.29) | 0.13 | 214 | 1.09 | (0.96 - 1.22) | 0.175 |
| Egger intercept | 246 | 0.001 | | 0.811 | 214 | -0.002 | | 0.444 |
| Q statistic | 246 | 527.1 | | $9.8\times10^{-23}$ | 214 | 188.1 | | 0.890 |
| | $FEV_1$/FVC and Adenocarcinoma | | | | | | | |
| | Manually filtered | | | | Manual filter and Outlier detection | | | |
| | $N_{SNPs}$ | OR | (95% CI) | P-value | $N_{SNPs}$ | OR | (95% CI) | P-value |
| IVW-RE | 247 | 1.19 | (1.01 - 1.40) | 0.043 | 228 | 1.24 | (1.08 - 1.43) | 0.002 |
| Maximum likelihood | 247 | 1.19 | (1.04 - 1.36) | 0.013 | 228 | 1.24 | (1.08 - 1.43) | 0.002 |
| Weighted median | 247 | 1.34 | (1.06 - 1.69) | 0.013 | 228 | 1.40 | (1.10 - 1.77) | 0.006 |
| MR RAPS | 247 | 1.20 | (1.02 - 1.42) | 0.029 | 228 | 1.28 | (1.10 - 1.48) | 0.001 |
| Egger intercept | 247 | -0.005 | | 0.182 | 228 | -0.006 | | 0.062 |
| Q statistic | 247 | 387.1 | | $3.7\times10^{-8}$ | 228 | 208.5 | | 0.806 |

IVW – RE : Inverse variance weighted – multiplicative random effects
RAPS : Robust adjusted profile score – Huber loss function

**Supplementary Figure 1:** Flow chart detailing the main quality control (QC) steps for the genome-wide association analyses of pulmonary phenotypes in the UK Biobank

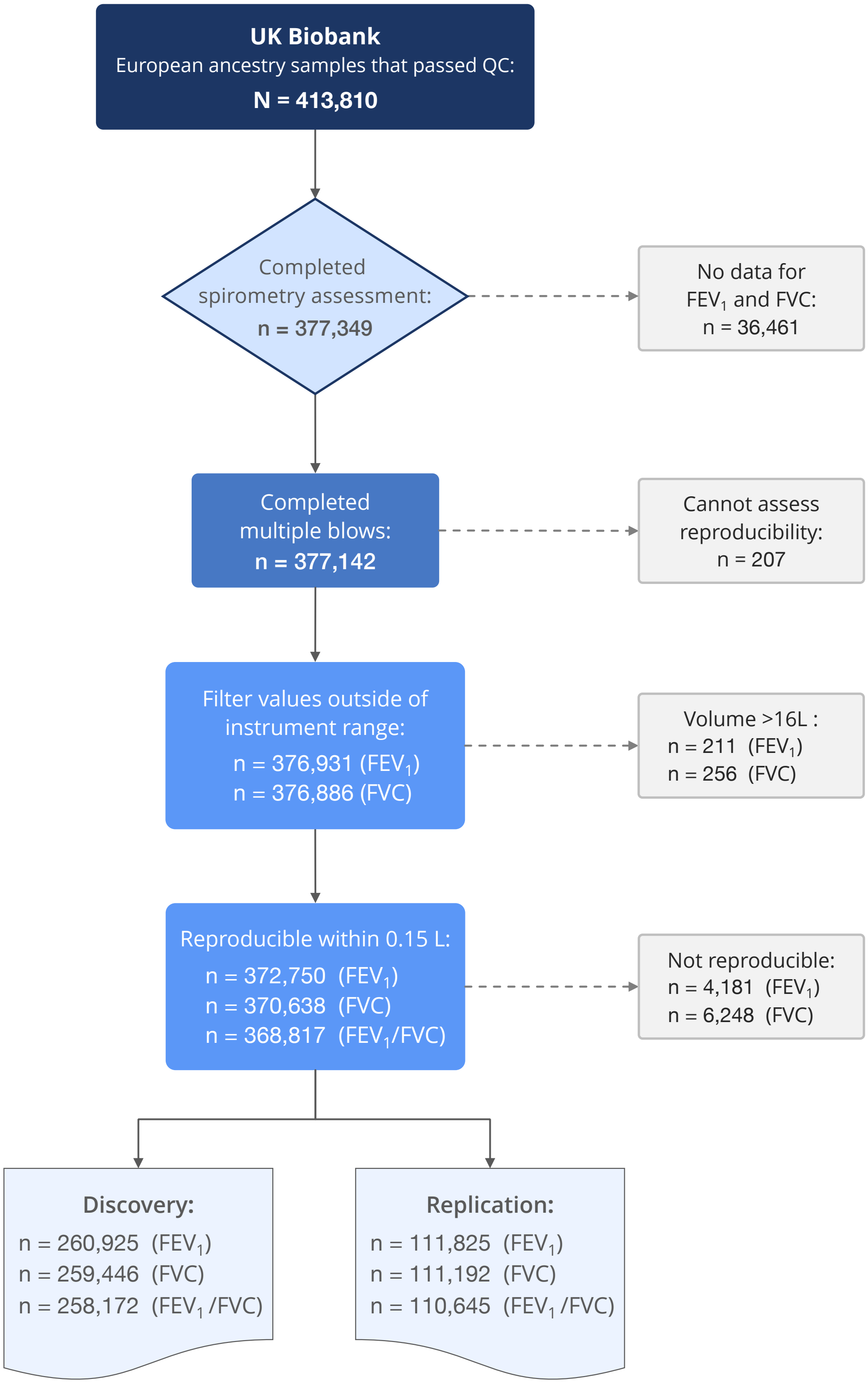

**Supplementary Figure 2:** Estimated power for Mendelian randomization analyses, based on the available sample size in the OncoArray dataset (29,266 lung cancers, 11,273 adenocarcinoma and 7,426 squamous cell carcinoma cases, and 56,450 controls), and assuming that the genetic instruments explain 5%, 3%, and 2% of variation in $FEV_1/FVC$, $FEV_1$, and FVC, respectively.

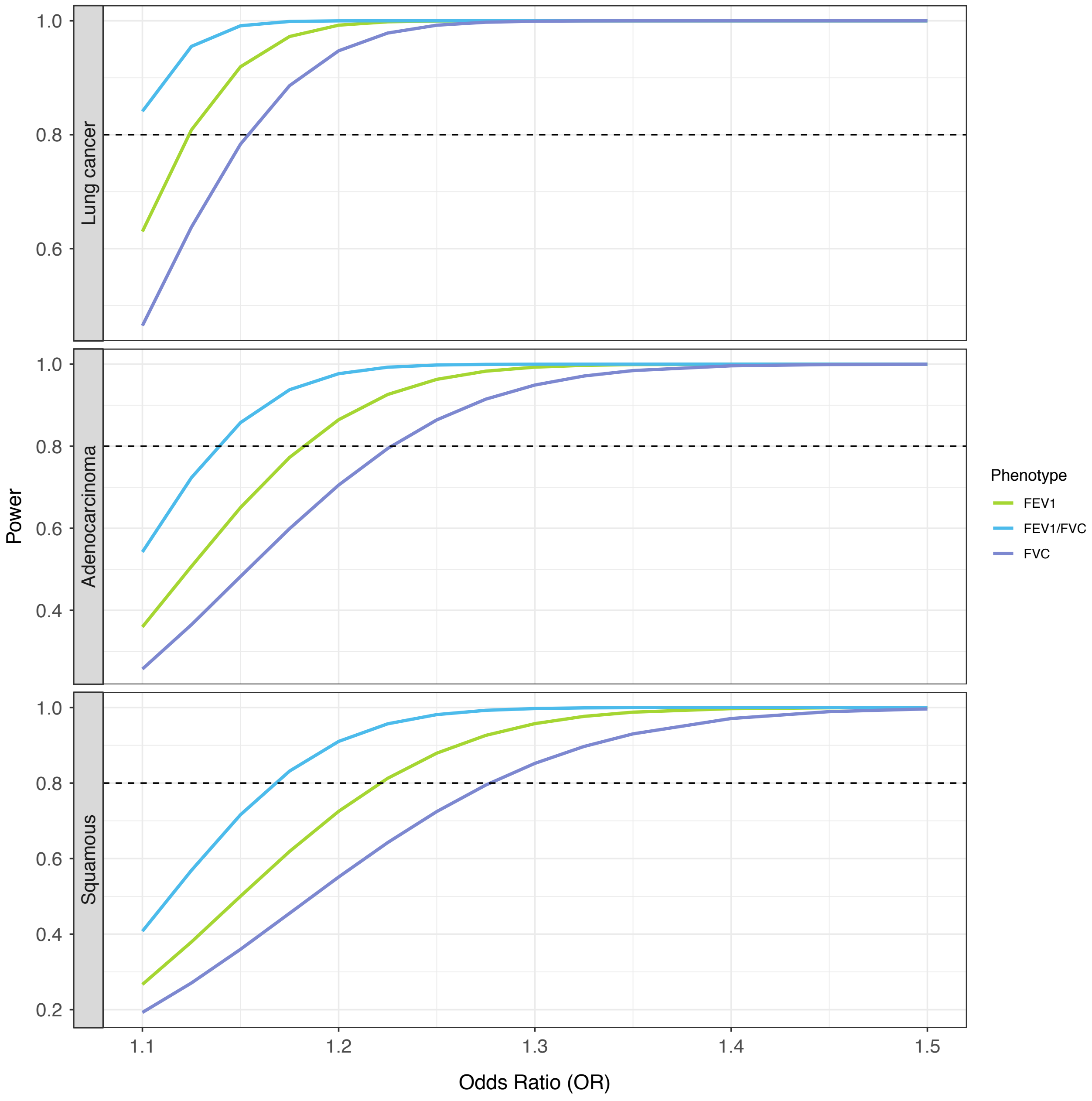

**Supplementary Figure 3:** Comparison of SNP effect sizes for pulmonary function genetic instruments estimated in the full UK Biobank cohort (original) and in ever smokers only (n=139,562 to 138,019) with adjustment for continuous cigarette pack-years rather than pack-year categories and years since quitting smoking (adjusted). Differences in the effect size distribution were tested using Wilcoxon rank sum test. All correlation coefficients were significant at $p<1\times10^{-40}$.

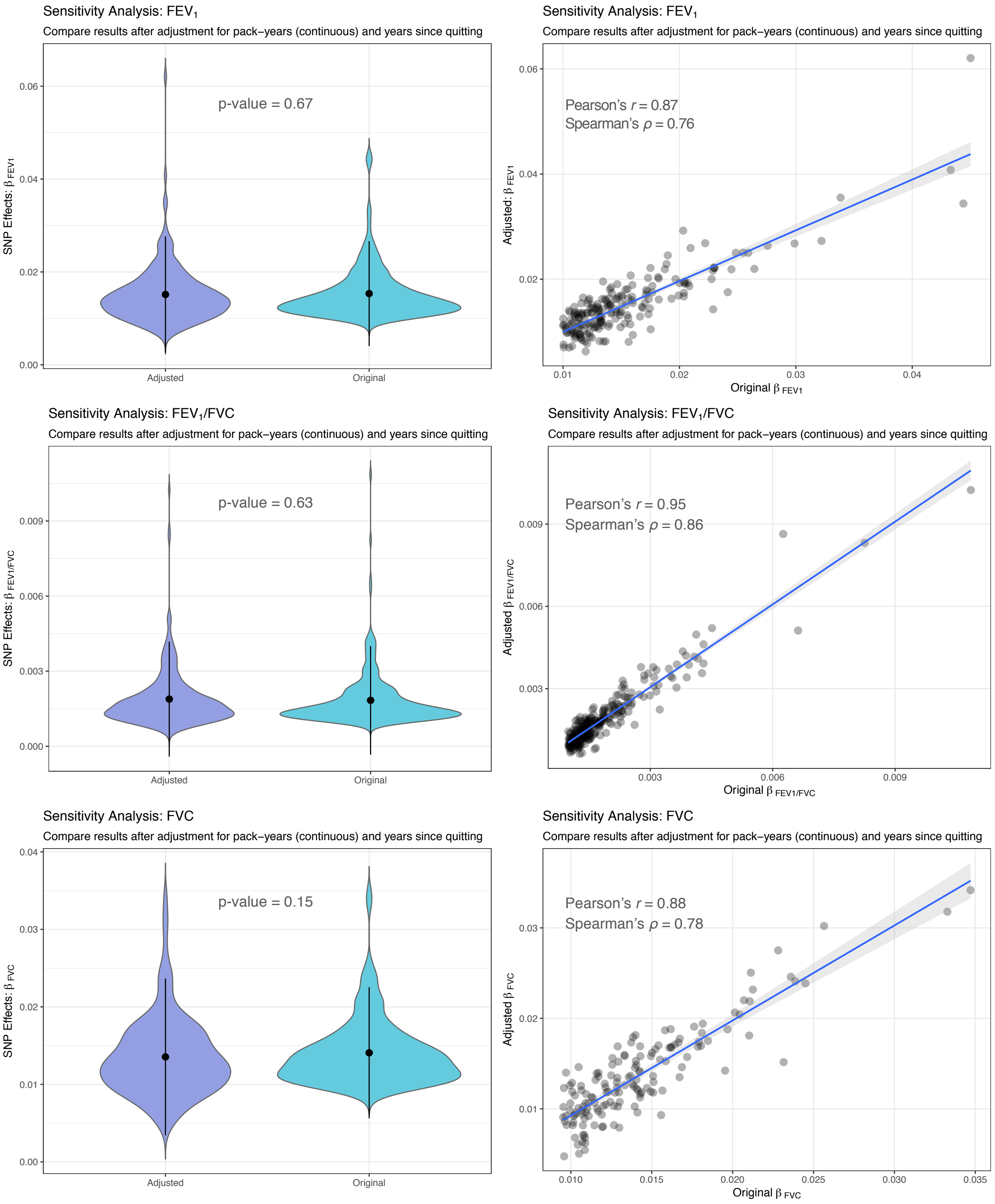

**Supplementary Figure 4:** Comparison of associations with chronic obstructive pulmonary disease (COPD), defined as $FEV_1/FVC<0.70$) for $FEV_1/FVC$ and $FEV_1$ genetic instruments. Effect estimates for COPD were estimated in the UK Biobank dataset using logistic regression with adjustment for age, sex, height, cigarette pack-years, genotyping array, and genetic ancestry principal components (PC1-PC15). A total of 54,711 cases and 313,640 controls were included in the analysis. The red solid line corresponds to genome-wide significance ($p=5\times10^{-8}$) and the red dotted line indicates nominal significance (p=0.05).

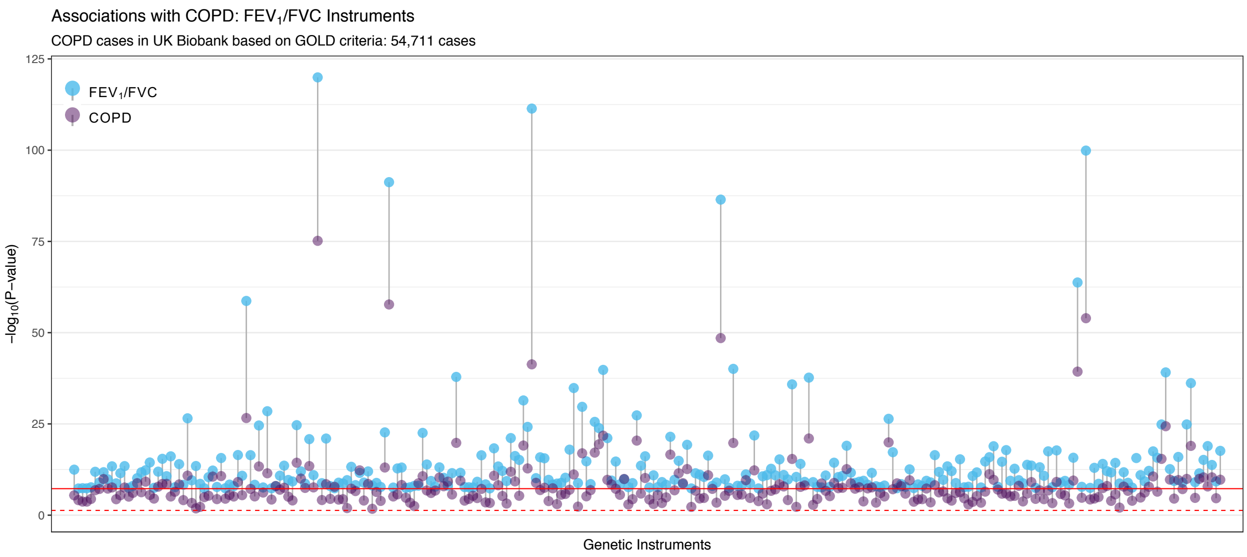


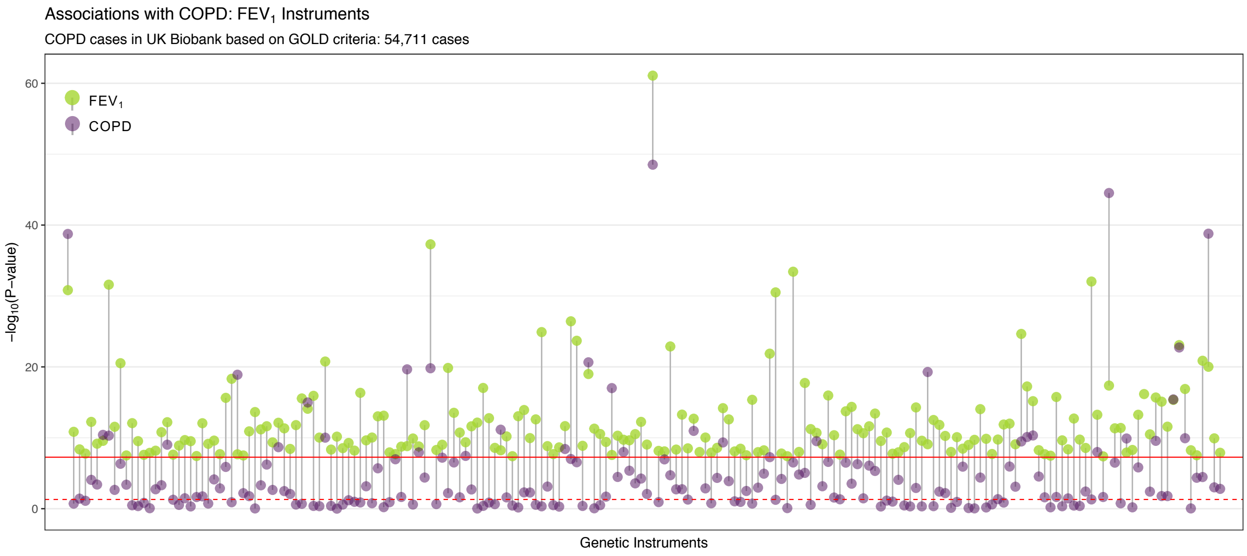

**Supplementary Figure 5:** Analyses of eQTL effects based on lung tissue gene expression data from the Laval Direction of eQTL effects was considered consistent if increased expression resulted in impaired pulmonary function and increased lung cancer risk (or increased $FEV_1$ or $FEV_1$/FVC and an inverse association with lung cancer). Lung cancer associations with $p<5\times10^{-4}$ were considered statistically significant and are labeled in the scatterplot. Odds ratios (OR) for lung cancer per unit increase in predicted gene expression, and corresponding associations with $FEV_1$ and $FEV_1$/FVC, are presented for all genes achieving p<0.05 in the lung cancer eQTL Mendelian randomization analysis.

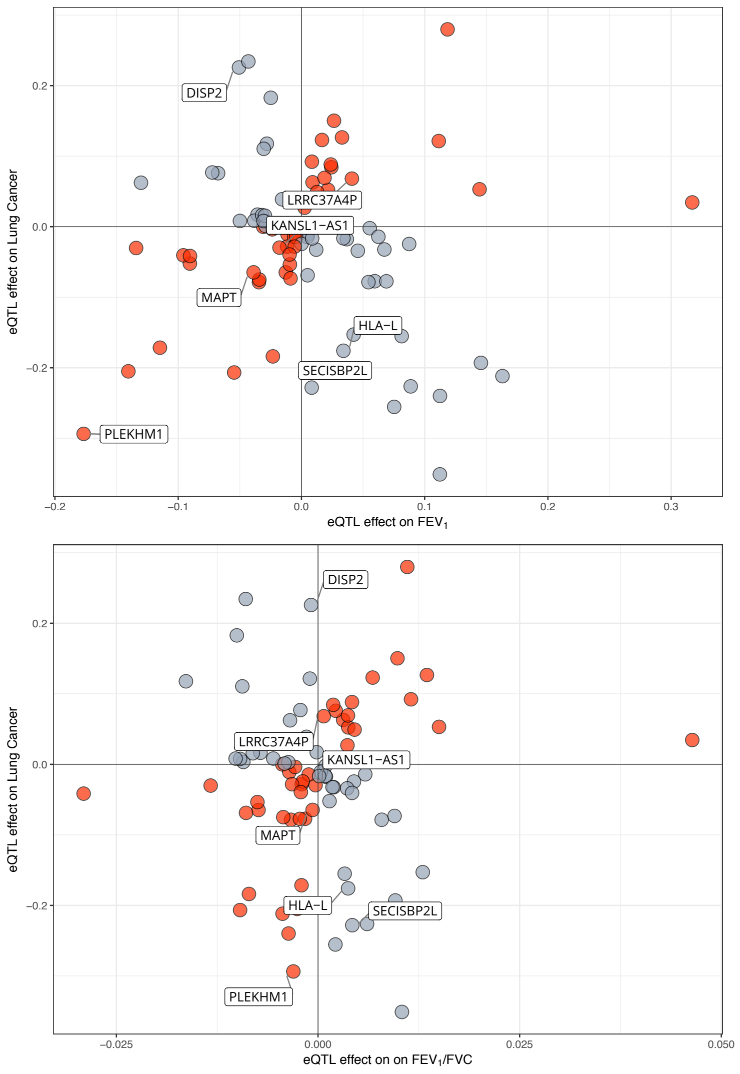


| Gene | OR | (95% CI) | $P_{Lung\ cancer}$ | $P_{FEV_1}$ | $P_{FEV_1/FVC}$ |
|---|---|---|---|---|---|
| *SECISBP2L* | 0.80 | (0.73-0.86) | $5.2\times10^{-8}$ | 0.191 | $3.8\times10^{-11}$ |
| *HLA-L* | 0.84 | (0.78-0.90) | $1.6\times10^{-6}$ | $4.0\times10^{-10}$ | $1.2\times10^{-11}$ |
| *DISP2* | 1.25 | (1.11-1.41) | $1.6\times10^{-4}$ | $3.9\times10^{-8}$ | 0.354 |
| *MAPT* | 0.94 | (0.90-0.97) | $4.6\times10^{-4}$ | $3.0\times10^{-62}$ | $4.3\times10^{-3}$ |
| *KANSL1-AS1* | 0.97 | (0.95-0.99) | $4.6\times10^{-4}$ | $3.0\times10^{-62}$ | $4.3\times10^{-3}$ |
| *LRRC37A4P* | 1.07 | (1.03-1.11) | $4.6\times10^{-4}$ | $3.0\times10^{-62}$ | $4.3\times10^{-3}$ |
| *PLEKHM1* | 0.75 | (0.63-0.88) | $4.7\times10^{-4}$ | $2.9\times10^{-62}$ | $4.3\times10^{-3}$ |
| *FBXL19* | 0.77 | (0.65-0.92) | $4.4\times10^{-3}$ | $2.9\times10^{-8}$ | 0.116 |
| *AZGP1* | 1.06 | (1.01-1.12) | 0.012 | 0.020 | $1.3\times10^{-15}$ |
| *TRIM4* | 0.83 | (0.72-0.96) | 0.012 | $4.9\times10^{-3}$ | $8.1\times10^{-25}$ |
| *WNT3* | 0.84 | (0.74-0.97) | 0.014 | $9.0\times10^{-33}$ | 0.035 |
| *RBM6* | 0.80 | (0.66-0.96) | 0.016 | $8.2\times10^{-10}$ | $3.1\times10^{-5}$ |
| *HLA-DPA1* | 1.09 | (1.01-1.18) | 0.023 | $7.9\times10^{-9}$ | $4.9\times10^{-24}$ |
| *HLA-DPB1* | 0.81 | (0.68-0.97) | 0.023 | $7.9\times10^{-9}$ | $4.9\times10^{-24}$ |
| *HSPA4* | 1.32 | (1.04-1.68) | 0.023 | $2.5\times10^{-10}$ | $4.9\times10^{-9}$ |
| *UQCR10* | 1.20 | (1.02-1.41) | 0.027 | 0.046 | $1.4\times10^{-15}$ |
| *HHIPL1* | 1.26 | (1.02-1.56) | 0.030 | $7.9\times10^{-3}$ | $4.5\times10^{-8}$ |
| *PDXDC2P* | 0.98 | (0.97-1.00) | 0.033 | $8.3\times10^{-25}$ | 0.113 |
| *ITGA2* | 0.92 | (0.86-0.99) | 0.034 | $1.6\times10^{-10}$ | $3.0\times10^{-16}$ |
| *UQCC1* | 0.79 | (0.63-0.99) | 0.039 | $4.2\times10^{-19}$ | $4.1\times10^{-3}$ |
| *MDM4* | 0.86 | (0.74-1.00) | 0.044 | $2.2\times10^{-11}$ | $7.1\times10^{-3}$ |

Histology-specific effects:

| Gene | OR | (95% CI) | $P_{Adeno}$ | Gene | OR | (95% CI) | $P_{Squam}$ |
|---|---|---|---|---|---|---|---|
| *SECISBP2L* | 0.64 | (0.57-0.72) | $3.1\times10^{-14}$ | *HLA-L* | 0.75 | (0.67-0.84) | $1.0\times10^{-6}$ |
| *PLEKHM1* | 0.64 | (0.51-0.80) | $8.8\times10^{-5}$ | *DISP2* | 1.30 | (1.08-1.57) | $6.2\times10^{-3}$ |
| *LRRC37A4P* | 1.11 | (1.05-1.17) | $8.9\times10^{-5}$ | *MDM4* | 0.74 | (0.58-0.95) | 0.017 |
| *KANSL1* | 0.96 | (0.93-0.98) | $8.9\times10^{-5}$ | *HLA-DRB6* | 1.06 | (1.01-1.11) | 0.022 |
| *MAPT* | 0.91 | (0.86-0.95) | $8.9\times10^{-5}$ | *STRA13* | 0.65 | (0.43-0.97) | 0.034 |
| *WNT3* | 0.74 | (0.61-0.88) | $1.1\times10^{-3}$ | *LOC285835* | 0.86 | (0.74-0.99) | 0.039 |
| *DISP2* | 1.21 | (1.03-1.42) | 0.021 | *MARCH3* | 0.89 | (0.79-1.00) | 0.048 |
| *MFAP2* | 0.87 | (0.77-0.98) | 0.026 | *C5orf63* | 0.76 | (0.58-1.00) | 0.048 |
| *HLA-L* | 0.90 | (0.81-0.99) | 0.034 | *SECISBP2L* | 1.05 | (0.92-1.20) | 0.44 |

**Supplementary Figure 6:** Visual summary of the Reactome pathway analysis depicting pathways that were significantly over-represented (FDR $p<0.05$) among genetic instruments for $FEV_1/FVC$ in never smokers. The size of circles (nodes) is proportional to the number of genes included in each pathway and the colors correspond to the q-value for each pathway.

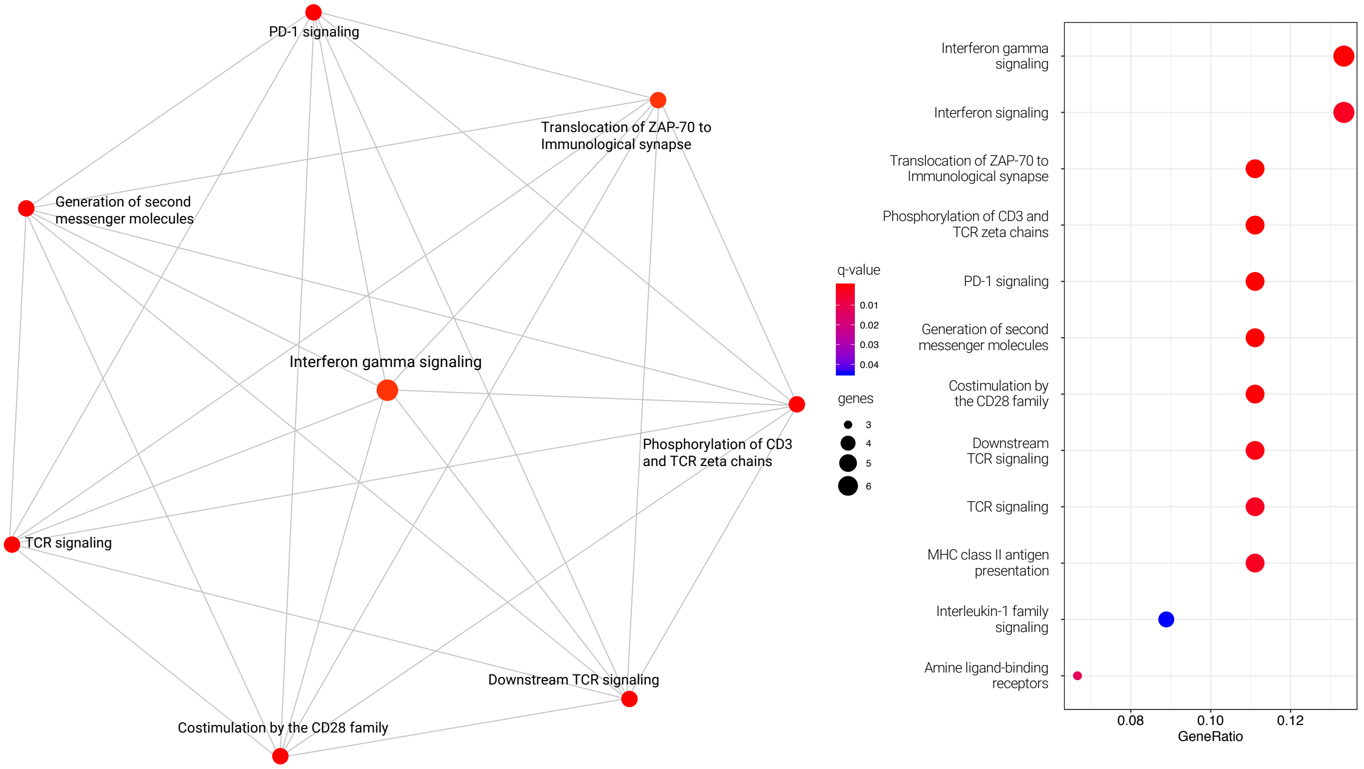

**Supplementary Figure 7:** Visual summary of the MSigDB pathway analysis depicting pathways that were significantly (FDR p<0.05) among genetic instruments for $FEV_1$ and $FEV_1/FVC$.

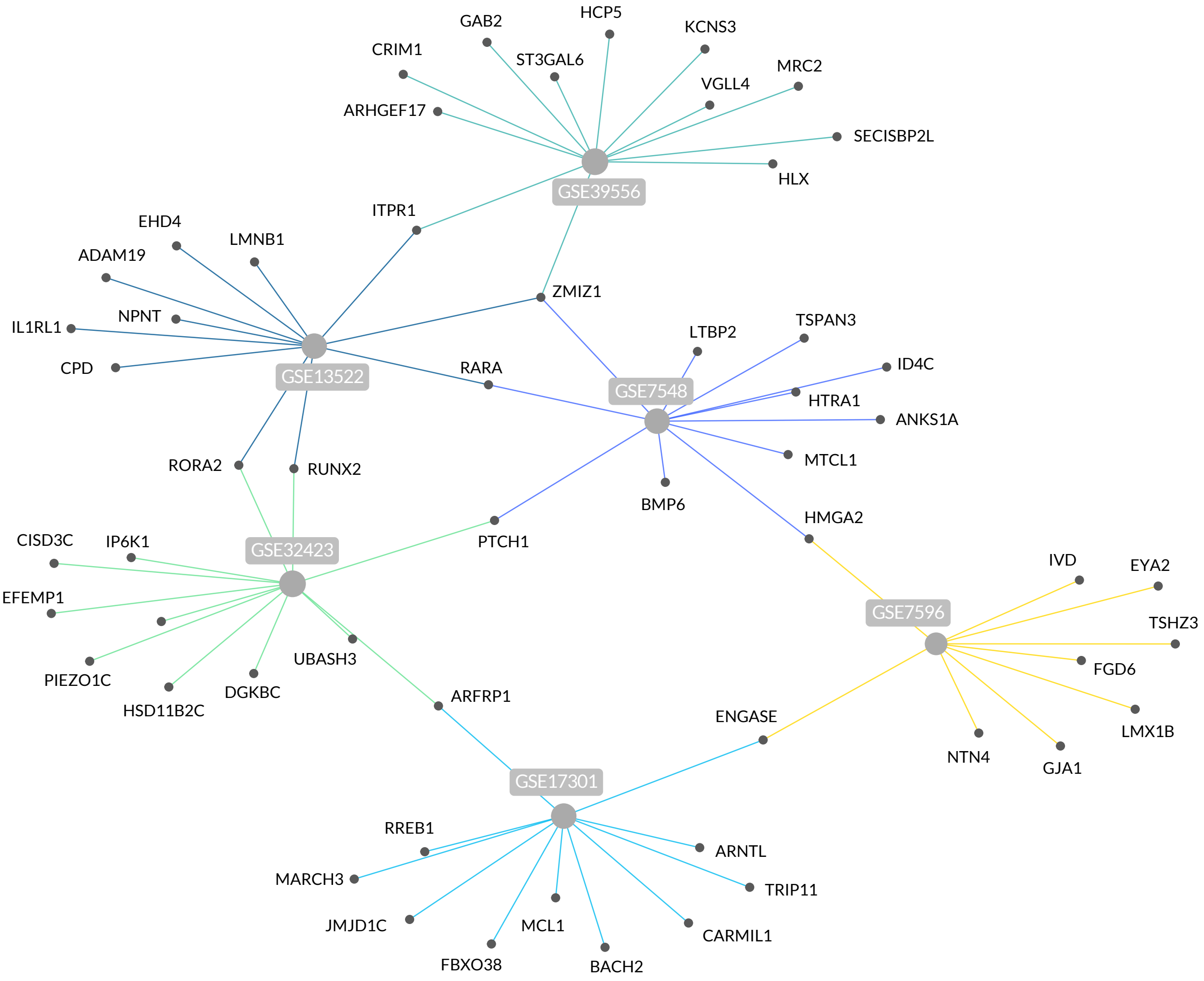


| ID | Description | p-value | q-value |
|---|---|---|---|
| GSE32423 | Up-regulated in memory CD8 T-cells treated with IL7 vs. IL4 and IL7 | $1.21\times10^{-5}$ | 0.024 |
| GSE39556 | Down-regulated in NK cells: untreated vs. pathogen-associated poly(IC) | $1.21\times10^{-5}$ | 0.024 |
| GSE13522 | Down-regulated during intradermal infection: wildtype (BALB/c) vs. IFNAR1 knockout | $1.91\times10^{-5}$ | 0.024 |
| GSE7596 | Down-regulated in T-cells in response to TGF-β: control vs. constantly active AKT1 | $2.16\times10^{-5}$ | 0.024 |
| GSE17301 | Up-regulated CD8 T-cells stimulated by INFA2 vs. IFNA5 | $6.19\times10^{-5}$ | 0.045 |
| GSE7548 | Up-regulated in lymph node CD4 T-cell: naïve vs. 28 days after immunization | $6.19\times10^{-5}$ | 0.045 |
